# Targets of SPEECHLESS and FAMA control guard cell division and expansion in the late stomatal lineage

**DOI:** 10.1101/2025.10.12.681868

**Authors:** Pablo González-Suárez, Yoëlle Hilbers, Alanta Budrys, Ao Liu, Sandra Richter, Dominique C Bergmann, Margot E Smit

## Abstract

Plant tissue development often relies on the specification of cell type initials with stem cell-like properties. These later undergo differentiation, losing division potential and acquiring specific identities and functions. In the stomatal lineage, protodermal cells develop into guard cells (GCs) through the action of bHLH transcription factors (TFs) SPEECHLESS (SPCH), MUTE and FAMA. Existing models support that these regulators act sequentially, but recent evidence indicates that SPCH expression and function are retained in late stomatal cells. Here, we combine transcriptomic and genetic approaches to define SPCH’s function during the late stomatal lineage. We show that relative levels and activities of SPCH and FAMA control GC division and expansion. Through cell type-specific TF induction and mRNA sequencing, we identify late-lineage targets of both TFs, and through genetic perturbation of these targets, we demonstrate that their precise temporal regulation is required for proper GC morphology and function. Our findings reveal a previously unrecognized role for SPCH in late stomatal development and support a revised model in which the functions of stomatal bHLHs are not strictly separated in time.

## Introduction

Three related basic helix-loop-helix (bHLH) transcription factors (TFs), SPEECHLESS (SPCH), MUTE and FAMA; regulate sequential cell state transitions in the stomatal lineage, each of which requires unique regulation of identity and division (Smit and Bergmann, 2023). To start the lineage, protodermal cells express SPCH, which promotes asymmetric cell divisions (ACDs) with the smaller daughter cell becoming a meristemoid and gaining early stomatal identity (Lau et al., 2014; MacAlister et al., 2007). Continued oriented divisions of meristemoids and their sister stomatal lineage ground cells (SLGCs) determine tissue patterning and stomatal density. Next, MUTE enforces commitment to stomatal fate and guard mother cell (GMC) identity (Pillitteri et al., 2007). Additionally, MUTE prevents further ACDs and instead slows down the cell cycle, promoting a single symmetric cell division (SCD) (Han et al., 2018; Han et al., 2022; Zuch et al., 2023). Finally, symmetrically-divided cells stop dividing and differentiate into guard cells (GCs) through the action of FAMA, which blocks further divisions (Ohashi-Ito and Bergmann, 2006).

While SPCH, MUTE and FAMA are homologs that depend on heterodimerization with the same partner bHLHs (Kanaoka et al., 2008), they are not interchangeable and have distinct interactors (Davies and Bergmann, 2014; Liu et al., 2024). Still, their bHLH domains are highly similar and, as a result, they recognize the same motifs and can bind to largely overlapping loci (Davies and Bergmann, 2014; Liu et al., 2024). Functional differences among the homologs have mainly been attributed to their unique, stage-specific expression. For instance, expressing FAMA or phosphovariants of SPCH in the *MUTE* expression domain can partially rescue *mute* (Davies and Bergmann, 2014; McKown et al., 2023). However, expression timing alone is not sufficient to recapitulate their specificity as those same SPCH phosphovariants are unable to complement FAMA’s function (Davies and Bergmann, 2014), suggesting that more complex regulatory dynamics are at play.

Expression of *MUTE* and *FAMA* is strictly limited to specific stomatal cells, with FAMA protein accumulating a few hours prior to the symmetric division of GMCs and then remaining present throughout GC differentiation (Zuch et al., 2023). In contrast, recent work has revealed that *SPCH* transcription extends from protodermal cells to later in the stomatal differentiation trajectory (Lopez-Anido et al., 2021). This was unexpected since the three bHLHs were thought to be temporally and functionally segregated. However, an early expression peak and early progression arrest in *spch* do not exclude a role for SPCH in the latter half of the lineage. SPCH’s extended role has been further confirmed using lines that exhibit lower SPCH levels at late stages. Rescuing *spch* mutants with SPCH transgenes whose expression does not persist into guard cells or targeting SPCH with artificial microRNAs in GMCs show results in cells that divert from stomatal fate and acquire pavement cell characteristics (Lopez-Anido et al., 2021). Still, it remains unclear what transcriptional programs SPCH directs in the late lineage and how these intersect with FAMA’s functions in regulating cell division and differentiation.

After the SCD of the GMC, FAMA prevents additional cell divisions by repressing expression of D-type cyclins (Han et al., 2018; Weimer et al., 2018) and through its partnership with RETINOBLASTOMA-RELATED (RBR) (Matos et al., 2014). Disrupting these pathways causes abnormal GC divisions which have different orientations and cell identity outcomes depending on how they are induced. For example, ectopic expression of *CYCD7;1* in GCs induces their symmetric transverse division while maintaining GC identity (Weimer et al., 2018). In contrast, extra divisions resulting from shorter *FAMA* expression, loss of the FAMA-RBR interaction, or reduction in SWI/SNF components or HISTONE ACETYLTRANSFERASE 1 (HAC1) often are asymmetric and result in re-initiation of the stomatal lineage with the formation of stomata-in-stomata (Lee et al., 2014; Liu et al., 2024; Weimer et al., 2018). Conversely, earlier expression of *FAMA* or blocking the SCD leads to the formation of single guard cells (Boudolf et al., 2004; Han et al., 2022; Simmons et al., 2019).

In addition, FAMA is necessary and sufficient for GC identity, and broad FAMA expression (*35S::FAMA*) appears to induce GC morphology and gene expression in other aerial epidermal cells (Ohashi-Ito and Bergmann, 2006). Earlier *FAMA* expression within the leaf stomatal lineage results in earlier differentiation, often of single GCs, though the ability of FAMA to do this depends on plant stage, with embryos being recalcitrant to FAMA-driven GC formation (Han et al., 2018; Smit et al., 2023). Thus, the timing of FAMA activity must be carefully regulated. Two of FAMA’s downstream targets, WASABI MAKER (WSB/ERF51) and STOMATAL CARPENTER 1 (SCAP1), are required for GC differentiation. Stomata of *wsb scap1* double mutants often arrest at young undifferentiated stages, failing to form pores (Shirakawa et al., 2025). SCAP1 has been associated with processes essential for GC maturation including pectin methylesterification and K^+^ transport. In accordance, ∼50% of stomatal complexes in *scap1* mutants have abnormal morphology with impaired GC integrity and stomatal opening (Negi et al., 2013). Pore formation coincides with pectic homogalacturonan (HG)-degrading enzymes and de-methyl-esterified HG accumulating at the pore site, at least in part facilitated by SCAP1-mediated activation of pectin methylesterases (Rui et al., 2019). The cuticle surrounding the pore forms a cuticular ledge whose opening depends on OCCLUSION OF STOMATAL PORE 1 (OSP1), a lipase, and FUSED OUTER CUTICULAR LEDGE1 (FOCL1), a proline-rich cell wall protein (Hunt et al., 2017; Tang et al., 2020). Beyond these factors, however, our understanding of the factors regulating GC maturation downstream of FAMA is limited.

Here, we investigate the distinct and shared roles of SPCH and FAMA in late stomatal development. Specifically, we address three main questions: (1) how does SPCH interact with FAMA’s functionality, (2) how does SPCH regulate GC division and morphology? and (3) which of the targets of SPCH and/or FAMA during late stomatal development regulate GC formation? Through genetic manipulation, we show that misexpression of *SPCH* in committed and maturing guard cells can induce ectopic symmetric divisions without reverting cells to a stem-cell-like identity. Using a cell-type specific RNA-seq approach, we identify downstream factors that regulate GC division and differentiation and demonstrate that these control stomatal morphology. Taken together, our findings shed new light on the complex regulation of late stomatal development and reveal some of the factors that must be robustly regulated.

## Results

### Misexpression of SPCH in the late stomatal lineage induces ectopic guard cell divisions

Recently, *SPCH* expression was shown to extend to later stages of the stomatal lineage, resulting in co-expression with *MUTE* and *FAMA* and contradicting our assumption that the functions of stomatal bHLH TFs are strictly temporally separated (Lopez-Anido et al., 2021). Interrogating additional scRNA-seq datasets (Kim et al., 2023; Zhang et al., 2021), we confirmed that co-expression of *SPCH* and *FAMA* occurs in late GMCs and early GCs with 3-28% of *FAMA*-expressing cells also expressing *SPCH* (**Figure 1A-B**, **Figure S1A-D**). Across independent experiments, it was apparent that *SPCH* expression in these cells is relatively low compared to that of *FAMA* (**Figure S1E**). Using translational reporters, we confirmed that SPCH and FAMA proteins are present simultaneously in some GMCs and GCs, with SPCH at lower levels than FAMA as cells progress towards GC identity (**Figure 1C-D**, **Figure S2**). This led us to question how SPCH may function at these later stages since, contrary to its well-known role in initiating ACDs (MacAlister et al., 2007; Smit and Bergmann, 2023), a potential involvement in the late lineage is less understood. Based on previous results that a reduction in SPCH from GMC stage onwards can lead to additional early lineage divisions and loss of stomatal fate (Lopez-Anido et al., 2021), we hypothesized that SPCH might be involved in controlling cell division programs and/or stomatal fate maintenance.

**Figure 1.**
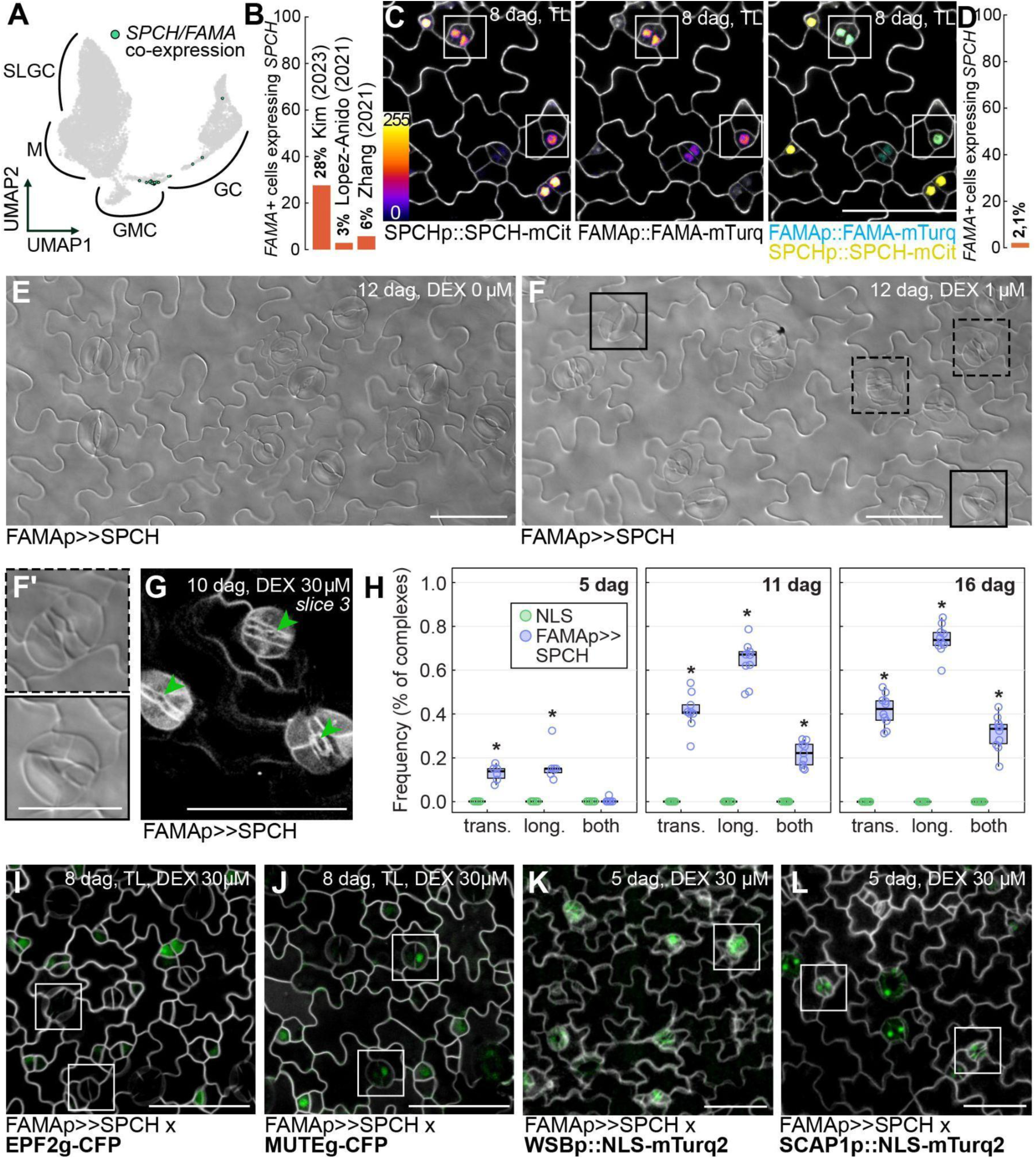
Misexpression of *SPCH* in the late stomatal lineage triggers abnormal guard cell divisions. **A.** UMAP plot based on scRNA-seq data from Lopez-Anido *et al*. (2021) highlighting cells that co-express *SPCH* and *FAMA* across stomatal development. SLGC: stomatal lineage ground cell, M: meristemoid, GMC: guard mother cell, GC: guard cell. **B.** Percentage of *FAMA+* cells that also express *SPCH* across three different scRNAseq studies. **C.** Representative confocal images showing expression of *SPCHp::SPCH-mCit* (left), *FAMAp::FAMA-mTurq* (middle) and merged (right) in true leaves of seedlings 8 days after germination (dag). The inset displays the colour legend for the fluorescence signal strength of *SPCH* and *FAMA*. Membranes are stained with propidium iodide (white). Scale bars indicate 50 µm. **D**. Percentage of cells containing FAMA-mTurq that also show SPCH-mCit across 2,550 FAMA+ nuclei of cotyledons imaged at 5 dag (N=17 leaves). **E-F.** Differential inference contrast (DIC) images of 12 dag *FAMAp>>SPCH* cotyledons grown with or without dexamethasone (DEX). Stomatal complexes with abnormal divisions are highlighted with solid squares (transverse divisions) or dashed squares (longitudinal divisions). Scale bars indicate 50 µm. **F’.** Close-up images of abnormal complexes. Scale bar indicates 25 µm. **G.** Confocal image showing the small pores in abnormal complexes upon *FAMAp>>SPCH* induction in 10 dag cotyledons. Membranes are visualized by combining the membrane marker *ML1p::mCherry-RCl2A* and propidium iodide staining (white). Scale bars indicate 50 µm. **H.** Quantification of additional divisions upon *FAMAp>>SPCH* or control *FAMAp>>NLS* induction in 5, 11 or 16 dag cotyledons (N=6-10 leaves). Complexes show additional longitudinal (long.), transverse (trans.) divisions, or both. **I-L.** Confocal images showing markers of meristemoids (*EPF2g-CFP*, **I**), GMCs (*MUTEg-CFP*, **J**) or guard cells (*WSBp::NLS-mTurq2*, **K**, and *SCAP1::NLS-mTurq2*, **L**) in 8 dag *FAMAp>>SPCH* cotyledons. Younger complexes that will undergo or have undergone additional divisions are highlighted using solid squares. Membranes are visualized using the membrane marker *ML1p::mCherry-RCl2A or* propidium iodide staining (white). Scale bars indicate 50 µm.

Having noted that expression and protein levels of FAMA are higher in these cells, we speculated that maintaining a low level of SPCH may be important for late lineage progression. To test this, we created a line where *SPCH* expression can be induced by dexamethasone (DEX) in the *FAMA* expression domain (i.e., late GMCs and GCs), FAMAp>>SPCH (OPp::SPCHg-Venus, FAMAp::LHG4-GR). Upon induction, ectopic SPCH resulted in additional GC divisions that were mostly longitudinal, leading to abnormal stomatal complexes reminiscent of the *fama* mutant (**Figure 1E-F**) (Ohashi-Ito and Bergmann, 2006). However, unlike in *fama* tumors, these did form pores, albeit small ones, similar to those reported in older *flp myb88* complexes (Lai et al., 2005; Yang and Sack, 1995). While we also observed transverse divisions, we found these mainly in mature complexes after the formation of completed pores (**Figure 1G-H**), suggesting that this became the dominant orientation when SCDs orientation was physically constrained by the presence of the pore. We observed these abnormal division phenotypes even after brief and transient induction of SPCH (**Figure S3**). Occasionally, we found instances of aborted stomatal complexes, which may indicate different effects of ectopic SPCH depending on the developmental cell status (**Figure S4**). Complexes with additional SPCH-driven divisions, however, did not form stomata-in-stomata. This contrasts with previous studies that destabilized GC fate by disrupting FAMA-RBR interactions (Matos et al., 2014) or by downregulation of chromatin remodelers (Liu et al., 2024). Thus, *SPCH* misexpression does not seem to induce re-initiation of the stomatal lineage and accordingly, *EPIDERMAL PATTERNING FACTOR 2* (*EPF2*), which is a direct target of SPCH during the early stomatal lineage, was not expressed in GCs upon *SPCH* induction (**Figure 1I**). Instead, *MUTE* was expressed in young GCs prior to additional cell divisions (**Figure 1J**), suggesting maintenance or repetition of GMC fate and associated programs. We found expression of the GC markers *WASABI MAKER/ ETHYLENE- RESPONSIVE ELEMENT BINDING FACTOR 51* (*WSB/ ERF51*) (Hachez et al., 2011; Shirakawa et al., 2025) and *STOMATAL CARPENTER 1* (*SCAP1*) (Negi et al., 2013) in SPCH-driven complexes (**Figure 1K-L**), confirming progression to GC identity.

### Guard cell division and morphology is controlled by levels and activity of SPCH and FAMA

FAMA’s role in promoting GC differentiation and inhibiting additional divisions has been described, (Hachez et al., 2011; Han et al., 2018; Matos et al., 2014; Ohashi-Ito and Bergmann, 2006) though the molecular functions of its targets and how they contribute to GC morphology have not been elucidated in detail. While increased levels of SPCH clearly affected GC division and morphology (**Figure 1F**), we found that *FAMA* induction neither promoted additional divisions on its own nor could abolish SPCH-driven divisions (**Figure S5**). This prompted us to ask whether SPCH and FAMA’s distinct effects during the later stages of the lineage could be at least partly explained by distinct regulatory abilities. Published ChIP-seq datasets suggest that their targets overlap with 73% of FAMA sites also bound by SPCH (**Figure S6**) (Liu et al., 2024). It is thus possible that differences in their contribution to GC development do not stem from specificity in DNA binding but rather from different co-factors resulting in either positive or negative regulation of specific target genes. To equalize and disrupt SPCH and FAMA functionally, we attached a SUPERMAN REPRESSION DOMAIN X (SRDX) tag (Hiratsu et al., 2003) to both, turning them into dominant repressors. Expressing SRDX-tagged SPCH and FAMA in the *FAMA* domain resulted in surprisingly similar phenotypes (**Figure 2A-D**). These included extra GC divisions (almost exclusively transverse); lobed GC morphologies, likely due to partial or complete transdifferentiation into pavement-like cells; and GC swelling, sometimes resulting in complex asymmetry. Taken together, the similar phenotypes that we observed supported the idea that misexpressed SPCH and FAMA largely find the same target genes but, under normal circumstances, have dissimilar effects on their expression.

**Figure 2:**
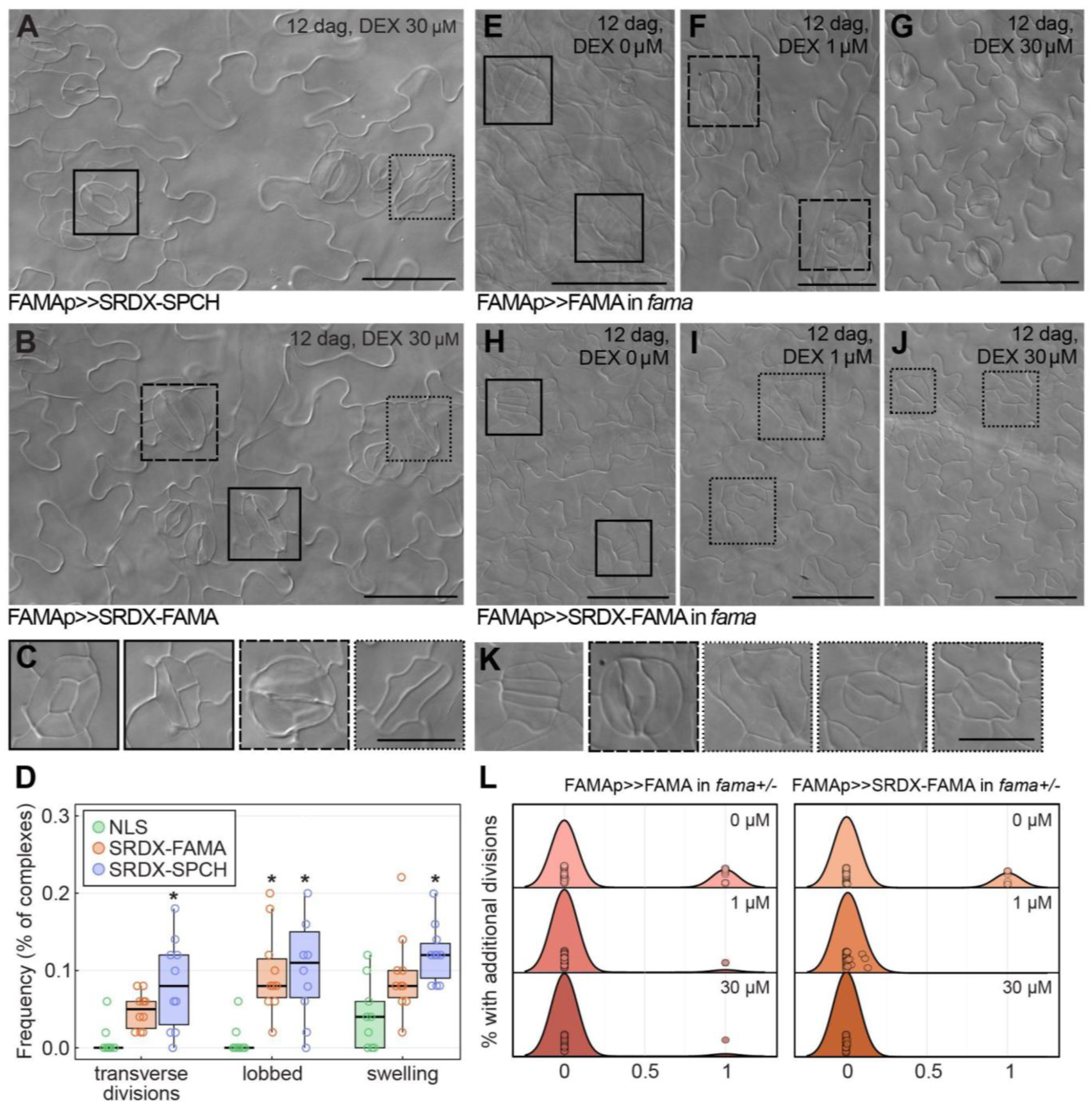
FAMA’s dual function in cell division and differentiation depends on dosage and transcriptional capabilities. **A-B.** DIC images of *FAMAp>>SRDX-SPCH* (**A**) and *FAMAp>>SRDX-FAMA* (**B**) cotyledons after 12 days of growth on DEX. Phenotypes are highlighted as follows: abnormal transverse divisions (solid squares), lobed guard cells (dotted squares) and guard cell swelling (dashed squares). Scale bars indicate 50 µm. **C.** Close-up images of guard cell morphologies observed in *FAMAp>>SRDX-SPCH* (**A**) and *FAMAp>>SRDX-FAMA* (**B**). Frame border represents the type of phenotype: extra transverse divisions (solid), lobbed guard cells (dotted) and swelling (dashed). Scale bar indicates 25 µm. **D.** Quantification of observed guard cells phenotypes upon SRDX-FAMA or SRDX-SPCH induction shown in A-C (N=10 leaves). Y-axis indicates the percentage of stomatal complexes that display specific phenotypes. Asterisks indicate statistical differences between each transgenic line and the *FAMAp>>NLS* control (ANOVA, Tukey HSD, p<0.05). **E-G.** DIC images of 12 dag cotyledons of *fama* with inducible *FAMAp>>FAMA* grown with DEX at 0 µM (**E**), 1 µM (**F**) and 30 µM (**G**). Solid squares indicate *fama* tumors and dashed squares indicate stomatal complexes with extra longitudinal divisions. Scale bars indicate 50 µm. **H-J.** DIC images of 12 dag cotyledons of *fama* with inducible *FAMAp>>SRDX-FAMA* grown with DEX at 0 µM (**H**), 1 µM (**I**) and 30 µM (**J**). Solid squares indicate *fama* tumors and dotted squares indicate stomatal complexes with abnormal morphology, including small or lobed guard cells, often with a crooked division plane. **K.** Representative close-up images of guard cell morphologies observed in *fama* mutants transformed with *FAMAp>>FAMA* (**E-G**) and *FAMAp>>SRDX-FAMA* (**H-J**). Scale bar indicates 25 µm. **L.** Fraction of stomata with additional longitudinal divisions in *fama+/-* mutants transformed with *FAMAp>>FAMA* (left) and *FAMAp>>SRDX-FAMA* (right) grown in control media or media supplemented with either 1 µM or 30 µM DEX (N=32-41 leaves).

FAMA acts as a positive regulator of many of its target genes (Hachez et al., 2011; Ohashi-Ito and Bergmann, 2006; Shirakawa et al., 2025). Our data demonstrate that SRDX-tagged FAMA is capable of blocking cell division in the late lineage, indicating that some of its downstream effects do not require gene activation, in line with its negative regulation of CDKB1;1 (Hachez et al., 2011). To further investigate this, we introduced both *FAMAp>>FAMA* and *FAMAp>>SRDX-FAMA* into the *fama* mutant. As expected, our regular induction levels (30 µM) fully complemented the *fama* mutant (**Figure 2E-L**, **Figure S7**). However, we found that lower induction of *FAMA* (1 µM), while able to fully restore GC differentiation, failed to completely prevent additional divisions, resulting in stomatal complexes with more than two GCs (**Figure 2F**). Similarly, SRDX-FAMA was also capable of preventing *fama* tumours (**Figure 2H-K**), confirming that FAMA’s native function as an inhibitor of cell division is at least partly mediated through transcriptional repression (Hachez et al., 2011; Weimer et al., 2018). However, division planes were often crooked and resulting GCs either became large and pavement cell-like or remained small, only rarely developing a pore, suggesting that final complex morphology and pore formation also depend on transcriptional activation. All together, these data are in support of a model in which dual roles in division and differentiation require different levels and activities of FAMA. In addition, our findings suggest that SPCH and FAMA can target some of the same downstream targets in maturing stomatal cells. It is tempting to hypothesize that higher levels of SPCH could act in part by competing for binding sites. However, additional experimental evidence is needed to support this hypothesis. In addition, competition with FAMA might not be the main mode of action of SPCH in the late lineage since co-inducing both results in the same complex morphologies as SPCH alone (**Figure S5B**).

### Identification of putative SPCH/FAMA targets in the late half of the stomatal lineage

From our observations of lines misexpressing *SPCH*, *FAMA* and their SRDX-tagged (i.e., dominant negative) versions, it was apparent that modifying the level and activity of SPCH and FAMA had strong effects on GC morphology and division. We sought to identify which genes downstream of SPCH and FAMA are responsible for their effects on GCs. Previous attempts to find targets of SPCH and FAMA involved broad induction over long timeframes (Hachez et al., 2011; Lau et al., 2014; Lau et al., 2018), which challenged the identification of direct targets. To improve resolution, we instead used our cell type-specific inducible lines to identify and compare rapidly induced target genes of SPCH and FAMA in GMCs and GCs. First, we treated 10-day-old seedlings with DEX for 2, 3 or 4 hours and used the fluorescence of the Venus-tagged TF variants driven by a *FAMA* promoter (or a nuclear-localized Venus control, *FAMAp>>NLS*) to isolate cells through fluorescence-activated cell sorting (FACS). The resulting cell populations include all cell types where the *FAMA* promoter is active: mainly GMCs and GCs, and a small portion of myrosin cells. Subsequently, we sequenced mRNA of FACS-sorted cells with the goal of profiling expression changes upon TF induction exclusively in these cell types (**Figure 3A**). The high specificity we aimed for resulted in a limited number of affected target genes which both strengthened and limited our follow up approaches (**Figure S8A-B**). Averaging transcript abundances per time point allowed us to distinguish broad differences more clearly (**Figure 3B**). However, using the NLS control as reference, we found that many differentially expressed genes (DEGs) down-regulated in either SPCH or FAMA were enriched in gene ontology (GO) terms related to DNA replication, mitosis and asymmetric cell division (**Figure S8C**). This was expected for FAMA, which inhibits division (Ohashi-Ito and Bergmann, 2006; Weimer et al., 2018) but not for SPCH given the phenotypes we previously observed (**Figure 1**). We reasoned that this could be due to a substantial difference in YFP signal strength, resulting in more cells with weak signal being included in the NLS sample and thus skewing the collected cell composition relative to the SPCH and FAMA samples (**Figure S8D**). As our main interest was in differential targets of SPCH and FAMA, we instead focused our analysis on genes that were significantly up- or down-regulated over time by induction of either TF or DEGs between the two. With this approach, we were able to identify 169 and 1,164 putative targets of SPCH and FAMA, respectively, as well as an additional 252 DEGs between FAMAp-driven SPCH and FAMA (**Figure 3C-D**, **Table S1**), adding up to a total of 1,552 unique DEGs. Among FAMA targets, down-regulated genes were enriched in terms related to ribosome biogenesis and protein translation, in agreement with FAMA’s known role in reinforcing the transition from proliferation to differentiation. We were unable to detect any statistically enriched GO terms among SPCH targets.

**Figure 3:**
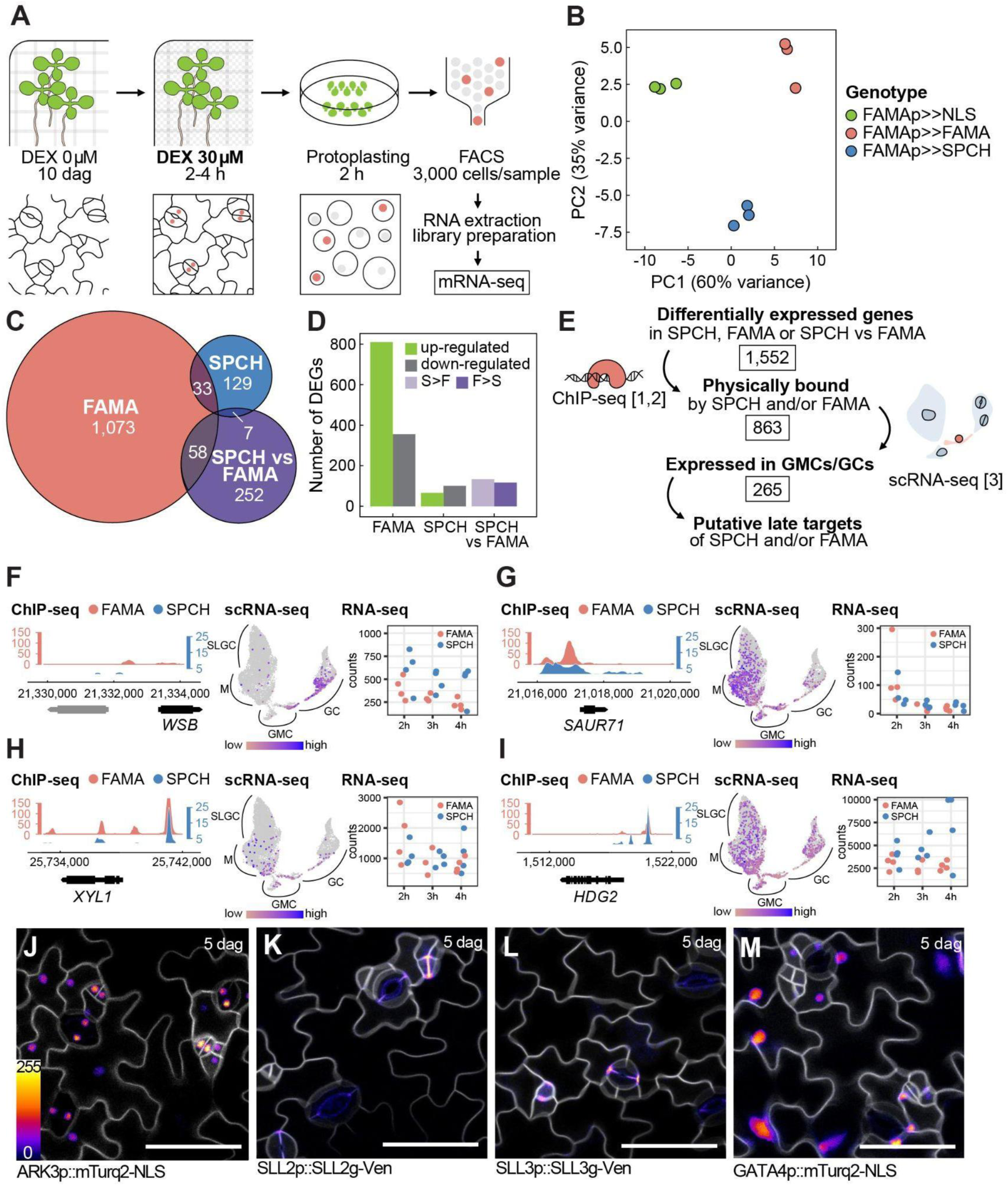
Identification of targets of SPCH and FAMA during the late stomatal lineage. **A.** Diagram illustrating the experimental approach followed for the identification of targets using inducible lines for SPCH, FAMA, and NLS (control) driven by the *FAMA* promoter (FAMAp>>X, i.e., *OPp::X-Venus*, *FAMAp::LHG4-GR*). Seedlings were initially grown for 10 days without DEX. After 2, 3, or 4 hours of DEX treatment, true leaves were collected and protoplasted. Late stomatal lineage cells were sorted using FACS to collect Venus+ cells marking the DEX-induced cells before library preparation and mRNA-seq. **B.** Principal component analysis (PCA) of mRNA-sequenced samples summarized per genotype. **C.** Venn diagram showing the total number of genes up- or down-regulated in *FAMAp>>FAMA* and *FAMAp>>SPCH* over time after DEX induction or differentially expressed between the two. **D.** Bar plot highlighting the number of genes either up- or down-regulated upon FAMA or SPCH induction, or differentially expressed when comparing the two. **E.** Schematic representation of the pipeline used for selecting putative targets of SPCH and/or FAMA in the late lineage for further analyses. **F-I**. Examples of the information used for our target gene selection for *WSB* (**F**), *SAUR71* (**G**), *XYL1* (**H**) and *HDG2* (**I**), including data from ChIP-seq (left) (Lau et al., 2014; Liu et al., 2024), scRNA-seq (middle) (Lopez-Anido et al., 2021) and our RNA-seq (right). Overviews for additional relevant genes used in this publication are in **Supplementary Figure S9**. **J-M.** Representative confocal images of transcriptional and translational reporters for newly identified late stomatal lineage markers and SPCH/FAMA target genes *ARK3* (**J**), *SLL2* (**K**), *SLL3* (**L**) and *GATA4* (**M**) showing their expression and/or localization in the late stomatal lineage in 5 dag cotyledons. The inset displays the colour legend for the fluorescence signal intensity. Cell membranes (white) are marked using the membrane marker *ML1p::mCherry-RCl2A*. All scale bars indicate 50 µm.

From the 1,552 unique DEGs, we sought to identify SPCH and FAMA targets that affect GC differentiation and morphology. Those could be either specifically upregulated in the late lineage or downregulated to prevent interference with maturation. As a first step towards the selection of candidate genes, we filtered for putative direct targets of SPCH and/or FAMA as indicated by promoter binding using published ChIP-seq datasets (Lau et al., 2014; Liu et al., 2024), which narrowed the list down to 863 genes (**Table S2**). Next, we made use of available scRNA-seq data (Lopez-Anido et al., 2021) to select 265 of these which were expressed in the stomatal lineage and/or with substantial transcriptional changes in the late lineage (**Table S3**). We surveyed these 265 genes considering their induction in our dataset, spatiotemporal expression according to scRNA-seq, induction by SPCH or FAMA in previous publications (Hachez et al., 2011; Lau et al., 2018), as well as their predicted function, to select 23 candidates for further study (**Figure 3E-I**, **Figure S9**, **Table S4**). We prioritized genes with predicted functions as transcription factors, receptors and other signaling factors or with known associated roles in cell division or expansion. Separately, we also selected some genes showing a sharp increase or decrease during the late stomatal lineage (see Materials and Methods).

Using transcriptional and translational reporters, we confirmed expression of most of the candidate genes in the stomatal lineage (**Figure 3J-M**, **Figure S10**). Expression patterns were already published for *SDD1* and *WSB* and our lines confirm expression in the late lineage (Hachez et al., 2011; Shirakawa et al., 2025; Von Groll et al., 2002) (**Figure S10C-D**). ARMADILLO REPEAT KINESIN 3 (ARK3) was previously identified as a SPCH target localized to the pre-prophase band during the ACD (Lau et al., 2014), but our transcription reporter indicates that its expression extends into the later stomatal lineage (**Figure 3J**). In addition, we identified three undescribed leucine-rich repeat family proteins as putative targets that are specifically expressed in the stomatal lineage, which we named STOMATAL LINEAGE LRRs 1-3 (SLL1-3) (**Table S4**, **Figure 3K-L**). Translational reporters revealed that SLL2 and SLL3 localize to cell membranes of GMCs and GCs. Finally, we also selected genes down-regulated by FAMA that were predicted to decrease in expression during stomatal differentiation, potentially having a negative effect on differentiation and, indeed, the GATA4 reporter showed that it is present in most epidermal cells but absent in GCs (**Figure 3M**).

### Putative targets of SPCH and FAMA in the late lineage control guard cell size and symmetry

Next, we asked what role these putative targets of SPCH and FAMA may have in symmetrically-dividing stomatal cells. In an effort to identify stage- and cell type-specific functions, we created misexpression lines for some of the selected genes using the *FAMA* promoter. Since most of these genes were down-regulated by FAMA (**Table S4**), we reasoned that ectopically increasing their levels would reveal their role during normal GC development while avoiding potential effects in the early lineage. In the case of TFs, we also used SRDX-fused versions to create a dominant negative effect. Each putative target was expressed as a fusion with a Venus tag, enabling us to confirm its expression in GMCs and GC as well as the subcellular localization of each gene’s product (**Figure 4A-B**, **Figure S11**). In the case of PDF1 (Abe et al., 1999; Abe et al., 2001), a putative extracellular proline-rich protein, we show for the first time that this protein is dramatically enriched at the cell wall or plasma membrane facing the stomatal pore (**Figure 4B**). To detect any effects from the misexpression of the putative targets on GC morphology and/or division, we measured stomatal size and density in 2-4 T2 lines per misexpression line in 12-day-old seedlings. We expected that changes in stomatal size would be indicative of altered GC morphology. In addition, stomatal density is inversely correlated to size, thus changes in size likely affect stomatal density as well (Doheny-Adams et al., 2012; Franks and Beerling, 2009). Based on this, we reasoned that both measurements would be informative initial readouts and criteria to select lines of interest for further experiments. Misexpression of several candidate genes did indeed affect stomatal size or density (**Figure 4C-D**, **Figure S12**). Specifically, *WSB* and *SMALL AUXIN UP RNA 71* (*SAUR71*) increased stomatal size while *ɑ-XYLOSIDASE 1* (*XYL1*) decreased it. In turn, misexpression of *XYL1* led to a large increase in stomatal density and *SRDX-GATA4* caused a slight decrease, opposite of *GATA4* (**Figure S12**).

**Figure 4:**
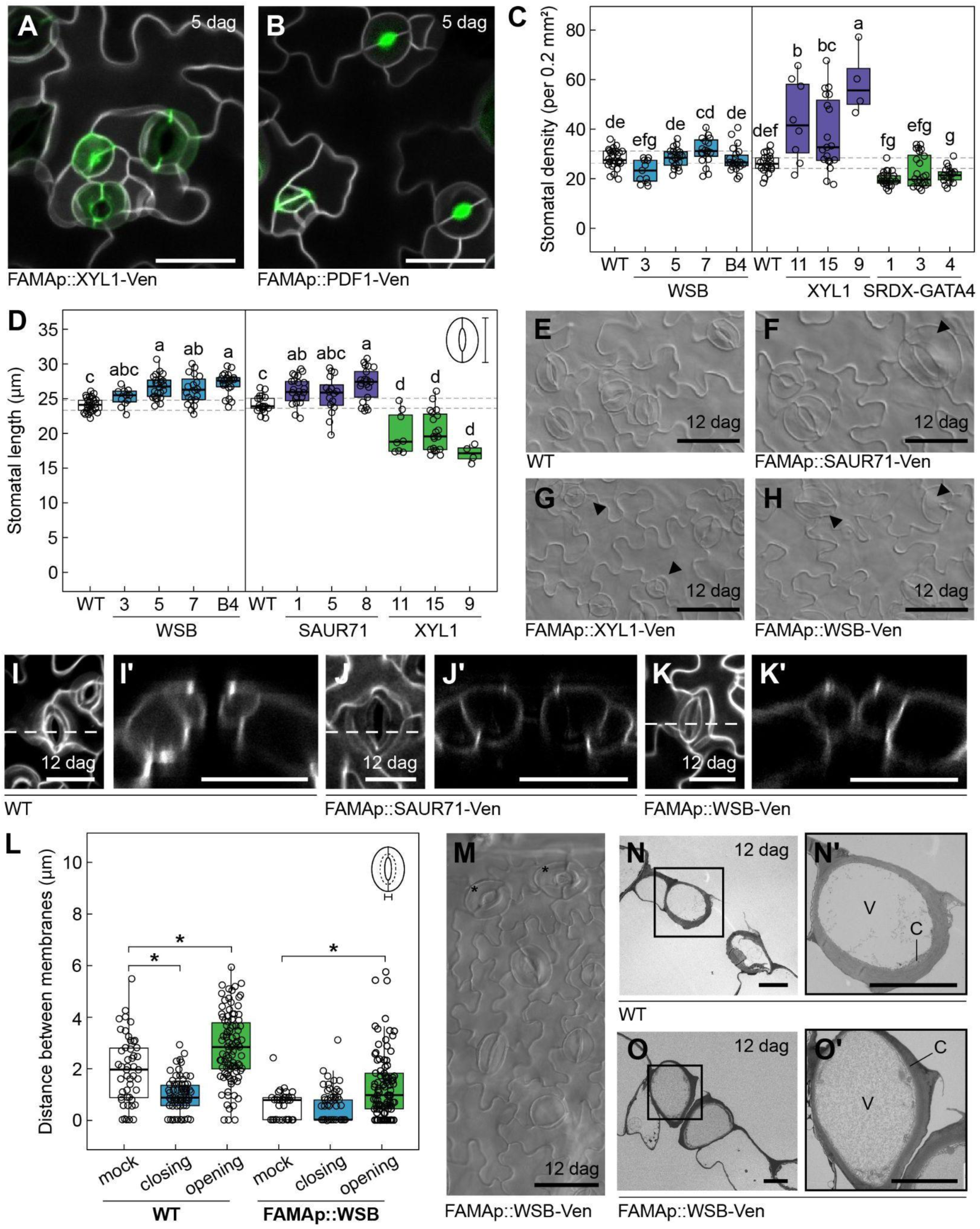
Targeted misexpression of targets of SPCH and FAMA in the late lineage impairs guard cell size and symmetry. **A-B.** Representative confocal images showing the subcellular localization of XYL1 and PDF1 upon misexpression driven by the *FAMA* promoter in cotyledons 5 days after germination (dag). Scale bars indicate 50 µm. **C-D.** Stomatal density (**C**) and length (**D**) in 12 dag cotyledons of T2 lines misexpressing putative targets of SPCH and FAMA (N=4-24 leaves). Dotted lines represent quartiles of wild-type (Col-0). Different letters indicate statistical differences between lines (ANOVA, Tukey HSD, p<0.05). **E-H.** DIC images of 12 dag cotyledons showing altered stomatal morphology in lines misexpressing *SAUR71* (GC bloating, **F**), *XYL1* (small asymmetric complexes, **G**) and *WSB* (large closed complexes and large single GCs, **H**). Arrowheads mark abnormal GC phenotypes, scale bars indicate 50 µm. **I-K.** Representative confocal images and cross-sections of lines that show bloated GCs or larger GC complexes, i.e., SAUR71 (**J**) and WSB (**K**), compared to WT (**I**). Dashed lines indicate the region corresponding to each accompanying cross section. **L.** Distance between cell membranes of sister GCs in complexes from 8 dag cotyledons of wild-type and *FAMAp::WSB* after treatment with mock, closing or opening solutions. Asterisks indicate statistical differences to wild-type (Student’s t test, p < 0.01) (N=3-5 leaves). **M.** DIC image showing open hydathode guard cell pores in 12 dag cotyledons of *WSB* misexpression lines, near the cotyledon tip, marked by asterisks. Scale bar indicates 50 µm. **N-O.** Cross-sectional TEM images of stomatal complexes in 12 dag cotyledons of wild-type (**N**) and *WSB* misexpression lines (**O**). V: vacuoles, C: cytosol. Solid squares indicate the regions corresponding to each accompanying close-up image. Scale bars indicate 5 µm. For all confocal images, membranes are marked with propidium iodide or the plasma membrane marker *ML1p::mCherry-RCl2A* (white).

A closer examination of the lines showing changes in size or density revealed two main classes of abnormal GC morphology: changes in GC size causing asymmetry between sisters and formation of large single GCs. For example, upon *SAUR71* misexpression, GCs were bloated to different degrees (**Figure 4E-F**). In these lines, pore morphology was normal, but the outer walls of the GCs were distended. In contrast, *XYL1* misexpression also resulted in asymmetrical GC complexes, but this was due to one sister GC failing to expand, resulting in either a very small pore or no visible pore at all (**Figure 4A**, **Figure 4G**). Lastly, misexpression of *WSB* resulted in an increase in stomatal size and obvious swelling of GCs. Unlike in *SAUR71*, the swelling was symmetrical between sister GCs and seemed to affect the pore, which appeared to be closed (**Figure 4H**). Cross sections through GCs of wild-type, *SAUR71* and *WSB* confirmed these observations, showing that complexes in lines misexpressing *SAUR71* remain open while those in lines misexpressing *WSB* have little to no space between the membranes of sister GCs (**Figure 4I-K**). We questioned whether these GCs were unable to open due to the bloating or because constitutive signaling kept them closed. To distinguish between these two possibilities, we treated cotyledons with stomatal opening and closing solutions (see Materials and Methods) and measured the distance between GC membranes at the pore. We found that GCs of *WSB* misexpression lines were able to open the pore (or at least increase distance between membranes) but at a reduced amplitude (**Figure 4L**, **Figure S13A-F**), suggesting that it is their swollen morphology that impairs their function. Interestingly, we noted that, in *FAMAp::WSB* leaves, the stomatal pores in the hydathodes (i.e., specific pores in the leaf margins associated to guttation) remain open, indicating that WSB might play a smaller role in their morphology (**Figure 4M**). Finally, we used transmission electron microscopy (TEM) to characterize the ultrastructural changes in *FAMAp::WSB* GCs that might result in the formation of closed complexes. Cell walls of *FAMAp::WSB* GCs looked similar to that of WT (**Figure 4N-O**, **Figure S13G-I**). However, there was increased electron density at the pore deposition site and in vacuoles, which leads us to hypothesize that a change in vacuole content might contribute to the GC swelling and inability to open.

### Timely expression of targets of SPCH and FAMA controls symmetric cell division

Misexpression of *WSB*, *SRDX-GATA4* and *SRDX-HDG2* in the *FAMA* domain resulted in large single GCs, occasionally with a kidney shape (**Figure 5A-C**, **Figure S14**). Optical cross sections of the single GCs revealed that they are often larger in all dimensions than typical 2-celled complexes (**Figure 5A’-C’**). Stomatal pore material, likely cutin and wax, accumulated in a ring-like structure centered in the exterior wall, although it appeared more diffuse than in wild-type (**Figure 5A-C, Figure 4I**). The accumulation of pore material did suggest that single GCs are undergoing stomatal differentiation. To confirm this, we crossed *FAMAp::WSB* with a transcriptional reporter for the mature GC marker *SCAP1* (Negi et al., 2013). Indeed, we detected SCAP1 in the single GCs induced by *WSB* misexpression, indicating that they display aspects of GC identity (**Figure 5D**).

**Fig 5:**
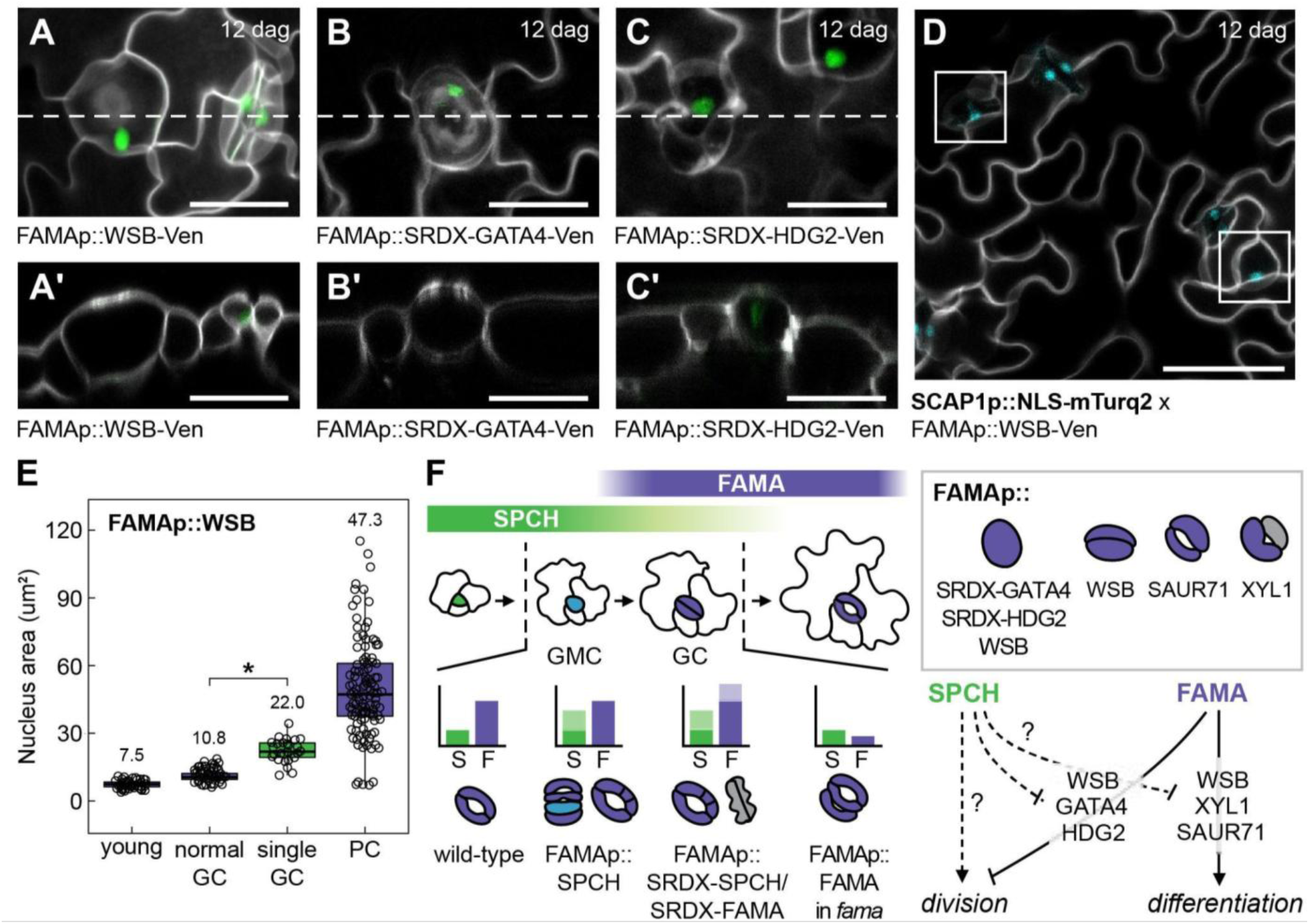
Misexpressed targets of SPCH and FAMA in the late lineage induce the formation of arrested single guard cells. **A-C.** Representative confocal images showing single GCs induced by *FAMAp*-driven misexpression of *WSB* (**A**) or SRDX-tagged dominant negative versions of *GATA4* (**B**) or *HDG2* (**C**) in 12 dag cotyledons. Dashed lines indicate the region corresponding to the accompanying cross sections, which emphasize the large volume of these single GCs in comparison to regular GCs and the accumulation of propidium iodide centrally on the outer surface of the cells. Scale bars indicate 25 µm. **D.** Representative confocal image highlighting the presence of a *SCAP1* transcriptional reporter in single GCs and paired GCs from 12-day-old cotyledons of *FAMAp::WSB.* Scale bar indicates 50 µm. Membranes are stained with propidium iodide and the plasma membrane marker *ML1p::mCherry-RCl2A* (white). **E.** Graph comparing nucleus area of different epidermal cell types in 8 dag cotyledons of *FAMAp::WSB* (N=2-5 leaves). Nuclei and membranes were stained with Hoechst and propidium iodide. Asterisks indicate statistical differences between normal and single GCs (Student’s t-test, p<0.05). **F.** Schematic overview summarizing results presented in this paper: the relations between SPCH, FAMA, their putative target genes, and the downstream GC features they affect (bottom right).

We were surprised to find that misexpression of putative SPCH/FAMA targets under the *FAMA* promoter was able to arrest the GMC division since *FAMA* expression starts late in the G2 phase of the cell cycle (Zuch-2023). Previous lines producing single GC phenotypes involved premature expression of FAMA using the *MUTE* promoter (Han et al., 2018) and plants with later increases in FAMA do not produce single GCs (**Figure 2**). Thus, it appears that, at the levels we induced, FAMA itself cannot prevent the stomatal SCD but several of its target genes can. To confirm at which stage of the cell cycle large single GCs arrest, we determined relative nuclear size by applying Hoechst staining and measuring different epidermal cell types. Nuclei of regular GCs were slightly larger than those of young stomatal cells whereas pavement cell nuclei appeared about 4-6 fold larger (**Figure 5E**, **Figure S15**). Nuclei of single GCs (SGCs) induced by WSB were ∼2-fold larger than those of regular GCs in the same leaves. In addition, they were of similar size as those formed upon the overexpression of a dominant negative version of CDKB1;1 (*CDKB1;1-N161*), whose nuclei were previously reported to be larger than wild-type due to arrest during G2 (Boudolf et al., 2004) (**Figure S15**). Therefore, our observations support the hypothesis that WSB-induced SGCs have differentiated during G2, similar to large SGCs formed upon misexpression of *SIAMESE- RELATED 1* (*SMR1*) or *CDKB1;1-N161* (Boudolf et al., 2004; Han et al., 2022). However, with our current data, it is unclear whether WSB, SRDX-GATA4 and SRDX-HDG2 promote the formation of SGCs only by blocking cell division, or whether they also actively promote premature GC differentiation (**Figure 5F**).

## Discussion

Classically, functional differences among SPCH, MUTE and FAMA have been attributed to a combination of distinct temporal expression and protein domains (Davies and Bergmann, 2014). Recent scRNA-seq data has challenged the temporal separation of all three, showing that while *SPCH* expression peaks in the early lineage, it persists in GMCs, and this late expression maintains stomatal fate commitment (Lopez-Anido et al., 2021). Here, we show that SPCH protein can be detected in GMCs and young GCs, resulting in these cells co-expressing SPCH and FAMA. Noting that SPCH levels are relatively low around the GMC to GC transition, we speculated that, while SPCH is required, its levels must be low for proper stomatal development. Indeed, inducing an excess of SPCH in the *FAMA* expression domain resulted in stomatal complexes with additional divisions and reduced GC size. Most abnormal divisions were longitudinal and resembled those reported for *fama* or *flp myb88* mutants (Lai et al., 2005; Ohashi-Ito and Bergmann, 2006), suggesting that increased SPCH in the late lineage either induces SCDs itself or impairs FAMA’s ability to inhibit them. As moderate late lineage overexpression of *SPCH* did not cause the same phenotype (Matos et al., 2014), high SPCH levels are apparently required for this change. Previous reports showed that ectopic GC divisions can be induced by short-lived FAMA, a disturbed FAMA-RBR interaction, or a reduction of chromatin remodeling factors (Liu et al., 2024; Matos et al., 2014). These mainly resulted in ACDs and stomatal lineage re-entry but could also result in occasional longitudinal divisions (Liu et al., 2024; Matos et al., 2014). Conversely, we found that excess SPCH did not reinitiate the lineage: it produced neither stomata-in-stomata nor expression of early stomatal markers. Instead, a persistence of *MUTE* indicates that GMC identity is maintained. Interestingly, this contradicts the idea that SPCH specifically promotes meristemoid ACDs, suggesting that the type of division induced by SPCH depends on the cellular context. Indeed, recent findings implicate SPCH in organ regeneration supporting the notion that SPCH may be employed to boost division capacity in multiple contexts (Tang et al., 2025). Thus SPCH’s function during GMC and GC stages likely depends both on precise dosage as well as the induction of cell type-specific target genes. Contrary to the effect of excess SPCH on GC division, we found that ectopically increasing FAMA did not have a substantial effect. This suggests that, while persistence of FAMA is key for GC maintenance, slight increases generally do not affect GC development, though a previous study reported that a FAMA transgene resulted in the formation of stomata-in-stomata indicating lineage re-entry (Lee et al., 2014).

To further disentangle SPCH and FAMA roles in GMCs and GCs, we equalized and disrupted SPCH and FAMA transcriptional regulation by tagging both with SRDX, turning them into dominant repressors (Hiratsu et al., 2003). Previously, FAMA-EAR was shown to produce *fama* (Ohashi-Ito and Bergmann, 2006), which is not completely in line with FAMA’s role as a repressor of some of its division-related targets (Hachez et al., 2011; Weimer et al., 2018). Here, we found that SRDX-SPCH and SRDX-FAMA resulted in similar phenotypes, including transverse divisions of GCs, complex asymmetry and lobed cell morphology suggestive of trans-differentiation to pavement cell identity. This indicates that both TFs target overlapping programs but, in native conditions, result in different outcomes. However, it does not reveal whether SPCH induction acts (in part) by impairing FAMA action through competition. By rescuing the *fama* mutant with low levels of FAMA, we found that low FAMA levels can indeed lead to additional symmetric divisions and GC complexes composed of three cells (**Figure 5F**). While this did not exactly replicate the *FAMAp>>SPCH* phenotype, it suggests that prevention of GC division requires low SPCH as well as high FAMA levels. In addition, it indicates that lower FAMA levels are needed to promote differentiation than to inhibit division. These two activities were previously partly unlinked in recent work as FAMA targets WSB and SCAP1 were found to specifically promote GC differentiation (Shirakawa et al., 2025). While in that work WSB was found to repress *SCAP1* in myrosin cells, we here show that WSB does not reduce SCAP1 levels in regular or single GCs (**Figure 5D**), highlighting differences in cell type-specific target gene regulation. Such differences are as-of-yet unexplained but might provide a starting point for future analysis of TF-target relationships across cell types. Finally, we also found that SRDX-FAMA was able to rescue *fama*’s GC division but not differentiation, indicating that the latter relies more strongly on FAMA acting as transcriptional activator. Alternatively, this could be the result of the SRDX domain rendering the FAMA protein non-functional, though its ability to prevent GC division indicates at least partial functionality.

Together, our experiments suggested that both SPCH and FAMA control GC development and that, despite similarities in their DNA binding (Lau et al., 2014; Liu et al., 2024), their respective effects are distinct. It remains unclear how SPCH and FAMA achieve their combined effects in GCs as no evidence for physical interaction or competition for binding sites has been found. SPCH was not recovered in proximity labeling experiments targeting FAMA (Mair et al., 2019), though this could be the result of lower SPCH abundance in GCs. Direct physical interaction between SPCH and FAMA might not occur if the two compete for binding sites when they regulate shared and unique target genes but the exact protein-level interactions and dynamics remain to be investigated. Regarding their functional outcomes, we show that SPCH induces cells to divide during both stomatal lineage initiation and when overexpressed in GMC/GCs, yet the types of divisions (ACDs vs. SCD) differ. Stage-specific chromatin landscapes and unique cofactors likely contribute to these different outcomes. We sought to understand which TF- and stage-specific target genes underlie these different outcomes. Using a cell type-specific transcriptomic approach and available ChIP-seq and RNA-seq datasets, we selected 23 putative targets. In investigating the function of these targets, we found that misexpression of several of them affected GC size and morphology leading to bloating, asymmetry between sister GCs and the formation of large single GCs. As some of these phenotypes overlap with those observed when inducing SPCH and SRDX-tagged SPCH or FAMA, these genes are likely to play a role in the execution of these programs.

Among the putative targets of SPCH and FAMA we identified several TFs. *WSB* was recently described as a key target of FAMA in GC differentiation (Shirakawa et al., 2025). Our misexpression of *WSB* resulted in overexpansion of GCs, which was accompanied by a permanently closed pore in most stomata, excluding hydathode guard cells, which remained open. These closed pores appeared to be the result of a change in physical cell properties rather than impaired signalling, as they were capable of opening and closing as measured by the distance between GC membranes albeit at a lower amplitude. Indeed, TEM revealed that, though cell walls in overexpanded GCs are similar to WT, their vacuoles are more electron-dense, suggesting an accumulation of unknown proteins. WSB was also linked to differentiation of myrosin cells (Shirakawa et al., 2025), whose vacuoles contain homogeneous electron-dense material essential for their role in defense (Andréasson et al., 2001; Shirakawa and Hara-Nishimura, 2018). The vacuolar density in bloated *WSB* GCs indicates that these cells might have some myrosin cells traits, resulting in a mixed cell state that interferes with GC function.

In addition to GC overexpansion, misexpression of *WSB* induced the formation of large SGCs that expressed *SCAP1* and formed a central deposition of pore-like material. Similar large SGCs have been previously reported when various regulators of cell division are impaired such as in *sol1 sol2 amiR-tso1* mutants (Simmons et al., 2019), CDKB1;1 dominant negative overexpression (Boudolf et al., 2004) and *SMR1* overexpression (Han et al., 2022). *SOL1* and its homologs are DREAM complex members that regulate cell fate and division (Simmons et al., 2019). The nuclear size of *WSB* SGCs indicated that they had undergone S-phase and differentiated during G2. We were surprised that SGCs could be induced by misexpression near the end of G2 considering that *FAMA* expression starts only ∼3 hours before the SCD (Zuch et al., 2023) and FAMA itself was only able to block division when expression under the *MUTE* promoter (Han et al., 2018). This indicates that the timing of FAMA targets is crucial for proper execution of the symmetric cell division. In addition to WSB, dominant negative SRDX fusions of GATA4 or HDG2 also resulted in the formation of SGCs. These findings indicate that FAMA directs the expression of several downstream TFs to inhibit divisions at the right time, with targets such as WSB providing a more direct path to division control. In a wild-type scenario, instead, timely activation of WSB and downregulation of GATA4 and HDG2 ensure that GCs do not divide any further after the first SCD.

From our putative targets, we also identified new stomatal marker genes and non-TF regulators of stomatal morphology. Three LRRs of unknown function, which we name *STOMATAL LINEAGE LRR* (*SLL1: AT3G17640, SLL2: AT3G20820, SLL3: AT5G23400*) are specifically expressed in the stomatal lineage with *SLL2* and *SLL3* acting as markers of the late stomatal lineage. SLL2, SLL3, the putative extracellular proline-rich protein PDF1, and the ɑ-xylosidase XYL1 all preferentially localize to GC cell membranes and/or pore structures. Misexpression of *XYL1* reduced GC expansion leading to smaller, sometimes asymmetric, complexes. XYL1 is required for xyloglucan maturation and has previously been shown to affect sepal elongation and trichome branching (Sampedro et al., 2010), suggesting a potential role for cell wall modifications during the late lineage in shaping GC morphology. XYL1 overexpression might decrease GC size by causing a premature or disproportional increased cell wall rigidity due to excessive xylose removal and higher xyloglucan cleavage, though this mechanism would require further investigation. In contrast, misexpression of *SAUR71* caused overexpansion of GCs, which sometimes showed blebbing but did form the pore. *SAUR71* was previously found in the stomatal lineage and, together with fellow members of the *SAUR41* subfamily, coordinates cell expansion and proliferation (Qiu et al., 2013). In light of our data, SAUR71 could contribute to GC expansion with its levels carefully constrained to control GC size.

## Conclusion

Our study shows that the final steps of stomatal development are regulated by FAMA and SPCH levels to ensure the formation of exactly two GCs of proper size and shape. We identify novel late stomatal genes downstream of SPCH and FAMA that control division and differentiation. We show that timely expression of these is critical for stomatal formation, and that their misexpression impairs final GC division and morphology (**Figure 5F**). Our data highlights that regulation of the single symmetric cell division preceding GC formation is tight with premature expression of WSB, one of FAMA’s targets, being able to block division where FAMA itself cannot. Extended expression of *SPCH* in the late lineage has been shown to be key for stomatal commitment (Lopez-Anido et al., 2021), but here we show that SPCH levels need to be kept low to prevent additional symmetric cell divisions, either directly or through competition with FAMA. This challenges the idea that SPCH specifically induces asymmetric divisions suggesting instead that the outcome of its division-promotive effect depends partly on cell identity. In addition, our data raises questions about the different effects of SPCH and FAMA on division and differentiation programs. While ChIP-seq has revealed a large overlap in bound genes, it remains to be discovered which of these are relevant and how differences on interactors enforce functional divergence between SPCH and FAMA as well as between early and late SPCH. Cell type-specific chromatin landscapes and proteomes could reveal driving factors in these differences.

### Study limitations

To identify targets of SPCH and FAMA specific to GMCs and GCs, we avoided broad misexpression to maintain normal tissue composition, instead choosing to use short, cell type-specific induction of SPCH or FAMA. While this allowed us to identify cell type- and stage-specific targets, the tradeoff was that we were only able to detect a limited number of differentially expressed genes. This was likely a result of noise resulting from induction and FACS approaches, inclusion of a small number of myrosin cells, and overall subtler changes at the whole transcriptome level leading to a higher number of false negatives. Although our strategy was successful in identifying several factors that act in GC development, future research studying highly specific developmental events may benefit from combining these specific approaches with more tractable and sensitive methodologies.

## Materials and Methods

### Plant material

All *Arabidopsis thaliana* mutants and transgenic lines used in this work, detailed in **Table 1**, are in the Col-0 background.

**Table 1.**
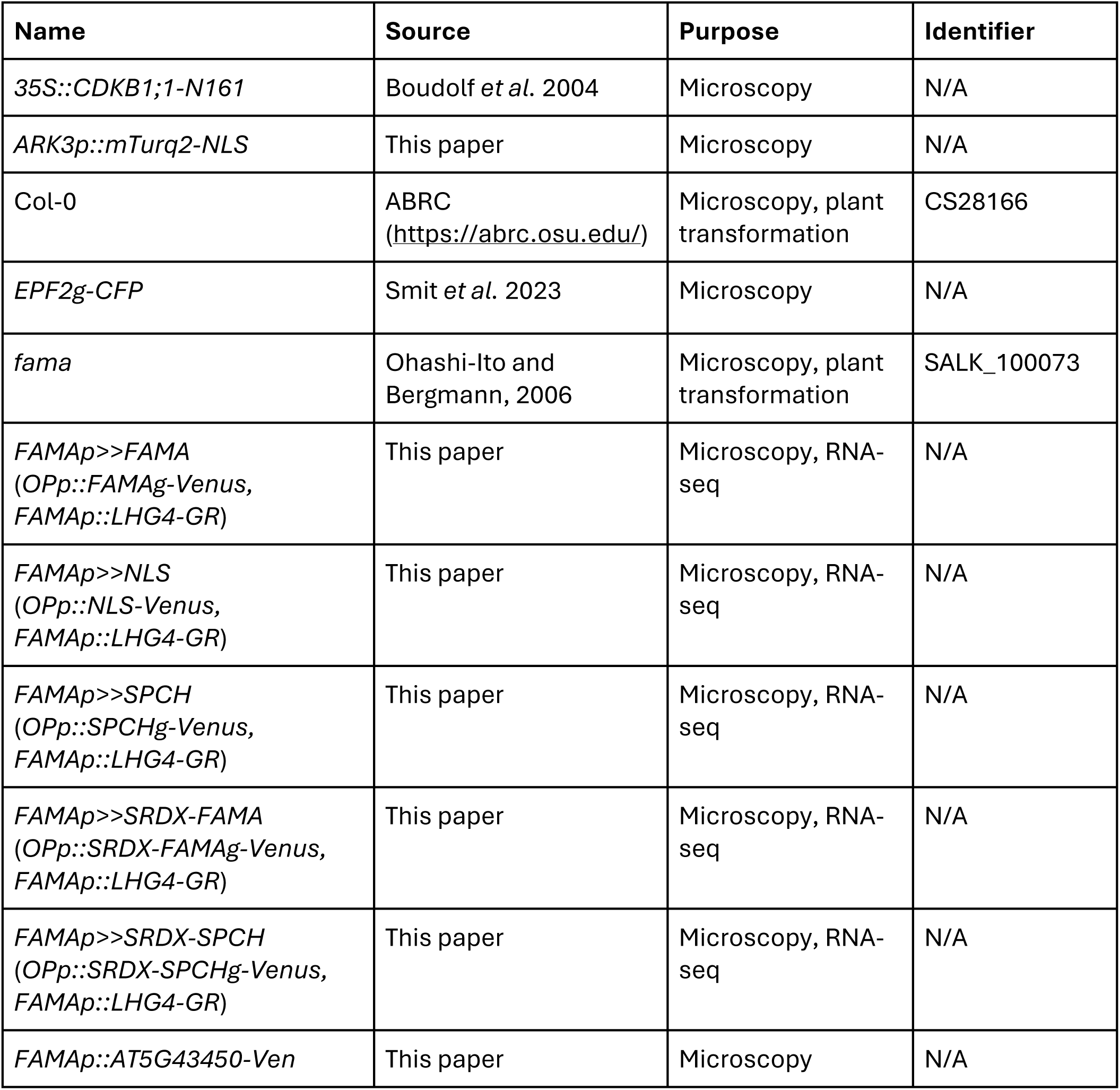
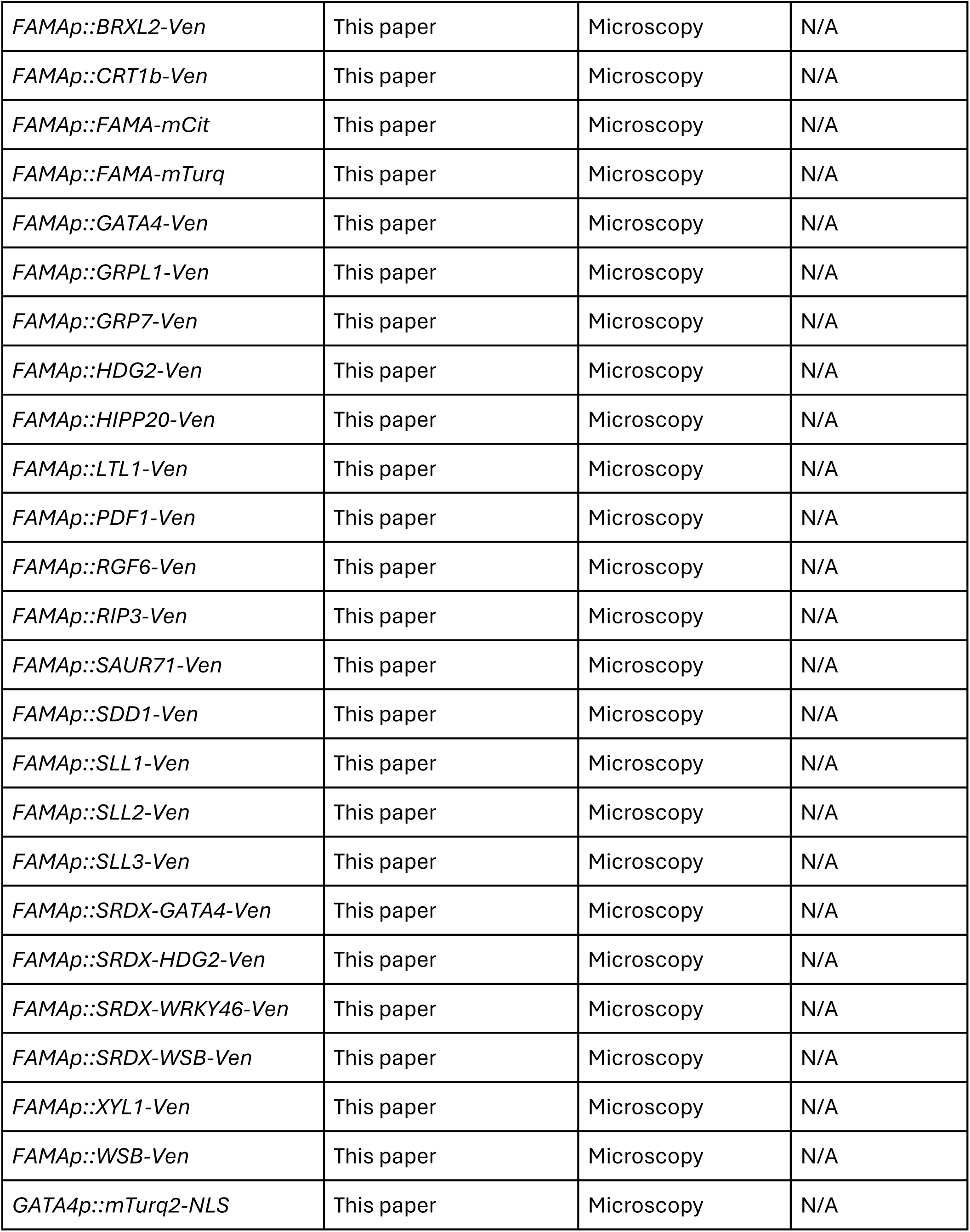
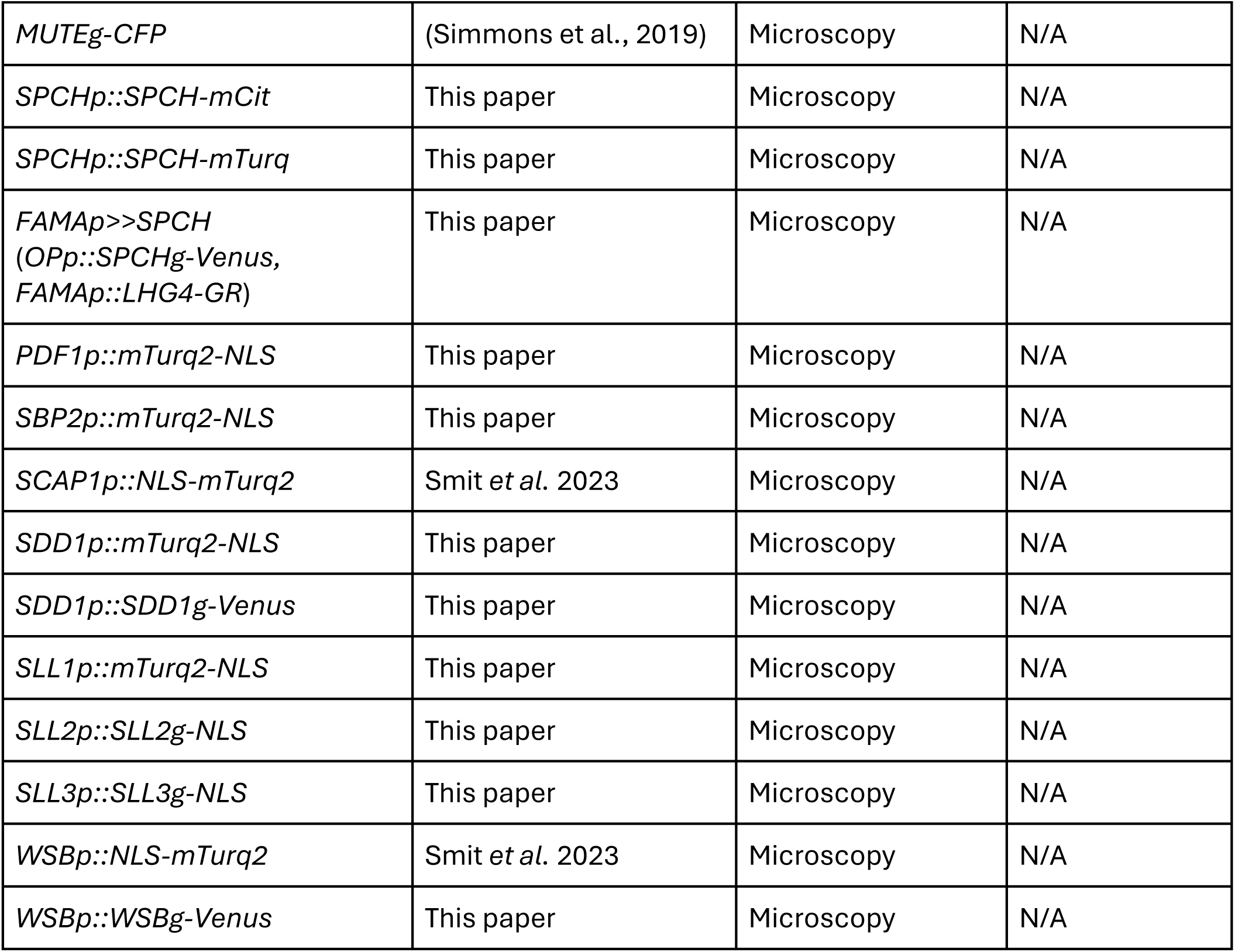
*Arabidopsis thaliana* mutants and transgenic lines used in this work.

### Plant growth conditions

Surface-sterilized seeds of *A. thaliana* were sown onto half-strength Murashige and Skoog (MS) growth media without sucrose and with 0.8% agar. Seeds were stratified for 2 days at 4°C, after which seedlings were grown for 5-12 days in a Percival growth chamber under standard long-day conditions (16h light: 8h darkness), light intensity of ∼120 µmol m^2^ s^-1^ and a day/night temperature cycle of 22°C/20°C.

### Molecular cloning and plant transformation

Primers used for cloning and genotyping are listed in **Table S5**. Constructs for plant transformation were cloned using GreenGate backbone pGreenII (Lampropoulos et al., 2013). Transgenic plants were generated by floral dip and transgenic seedlings were selected on ½ MS without sucrose based on antibiotic resistance (15 mg/L phosphothricin or 7.5 mg/L sulfadiazine).

### Microscopy

Abaxial leaf surfaces were imaged for all experiments. Differential inference contrast (DIC) images were taken with a Leica DMi8 inverted scope using 20x and 40x objectives and the tiling function. Fluorescence imaging was performed on a Leica SP8 or Stellaris confocal microscope with HyD and PMT detectors using 20x and 25x water objectives. Cell outlines were visualized using the plasma membrane marker *ML1p::mCherry-RCl2A* (Davies and Bergmann, 2014) or propidium iodide (10 μg/ml). All microscopy images were analyzed using FIJI. For fluorescent reporters, raw Z-stacks were projected with the ‘Sum Slices’ function.

### Stomatal size and density quantification

Stomatal size and density were measured from DIC images of cleared leaves mounted using Hoyer’s solution (gum arabic, glycerol, and chloral hydrate in water). For the quantification of stomatal density, all stomata in a set area were counted. For the quantification of stomatal size, the length of the first 25 stomata on a leaf were measured.

### Pore opening and closing assays

Opening and closing assays were performed using previously established protocols (Yu and Assmann, 2016). 8 dag seedlings were incubated in liquid ½ MS without sucrose supplemented with 50 mM NaCl (opening) or 100 µM ABA (closing). Seedlings were incubated for 2 hours in regular growth conditions (mock, opening) or in the dark (closing). The maximum distance between either membranes or pore depositions was measured from z-stacks where membrane and pore deposition was visualized using propidium iodide.

### DEX treatment

For longer treatments, seedlings were grown on plates containing 0, 1 or 30 µM dexamethasone (DEX). Short induction for RNAseq was performed by first growing seedlings on a nylon mesh on regular ½ MS without sucrose, then treating with liquid ½ MS without sucrose supplemented with 30 µM DEX on the plate for 2 minutes and finally transferring the mesh with seedlings to a fresh 30 µM DEX plate for 2-4 hours of additional growth.

### mRNA sequencing

GR-tagged induction lines were used for cell type-specific mRNA-seq. All 5 lines contained the *FAMA* promoter driving GR-LhG4, promoting expression of Venus-tagged *FAMA*, *SPCH*, *SRDX-FAMA*, *SRDX-SPCH* or a nuclear localization signal (*NLS*) via OPp (see Plant material). 10 dag seedlings of each line were treated with DEX to induce transgene expression. Upon induction, cells were protoplasted for 2 hours and used for fluorescence-activated cell sorting (FACS) to sort for Venus signal indicating successful induction. Separate samples of each line were collected after 2, 3 and 4 hours of DEX treatment. For each time point, a total of 3-4 biological replicates containing ∼3,000 cells each were collected. RNA was isolated with the Qiagen RNeasy Micro Kit (74004) and sequencing libraries were prepared with the SMART-Seq® v4 Ultra® Low Input RNA Kit from Takara following the manufacturer’s guidelines. Libraries were sequenced 50 bp with single end on an Illumina HiSeq 2000.

### mRNA-seq analysis

Raw reads were aligned to the *Arabidopsis thaliana* TAIR10 genome assembly using STAR and read counts were quantified by HTseq. Read counts were further analyzed using DESeq2 (Love et al., 2014) using the following pipeline. Initially, principal component analysis (PCA) was used for exploratory data analysis. To minimize data variability due to small time steps between samples, read counts were summarized per time point. For each genotype, differential gene expression was initially performed using *FAMAp>>NLS* as a control and testing for statistical significance of the genotype:time interaction with a likelihood ratio test following the developers’ guidelines. Gene ontology (GO) enrichment was performed on the lists of differentially expressed genes using clusterProfiler (Yu et al., 2012).

A second set of differential expression tests was performed using unsummarized read counts of *FAMAp>>FAMA* and *FAMAp>>SPCH*. For each genotype, three pairwise Wald tests were used to test for a significant effect of time on gene expression: (1) 2 h vs. 3 h, (2) 4 h vs. 3 h and (3) 4 h vs. 2 h. In addition, three additional pairwise Wald tests were used to test the effect of genotype within each time point comparing SPCH and FAMA samples.

The following approach was followed to select genes of interest for follow-up analyses (**Figure 3E**). The starting point was a list of unique differentially expressed genes (DEGs) identified from any of the six statistical comparisons, which added up to a total of 1,552 (**Table S1**). First, published ChIP-seq datasets for SPCH (Lau et al., 2014) and FAMA (Liu et al., 2024) were used to select DEGs whose genomic loci are bound by either transcription factor. Next, for the 863 remaining genes (**Table S2**), scRNA-seq plots were generated using available datasets (Lopez-Anido et al., 2021) and used to score spatiotemporal gene expression. Each of the 863 genes was scored with one of the following categories based on its preferential expression in specific cell types: (1) ‘young guard cell’, (2) ‘young other’, (3) ‘old guard cell’, (4) ‘stomatal’, (5) ‘differentiating’, (6) ‘early stomatal’, (7) ‘pavement cell’, (8) ‘broad’, (9) ‘sparse’, (10) ‘absent’, (11) ‘other’ or (12) ‘ubiquitous’. All genes with any of the first 6 categories (a total of 265, **Table S3**) were kept for further analysis while the rest were discarded. In the final selection step, we considered additional published datasets as well as functional predictions. Previous *SPCH* (Lau et al., 2014; Lau et al., 2018)(Lau-2014, Lau-2018) and *FAMA* (Hachez et al., 2011) induction datasets were used to determine which targets were shared across experiments. Further, we used the scRNA-seq plots to look closely at previously reported dynamics in the late lineage. Finally, we considered predicted gene functions to select genes that might affect GC morphology. Altogether, to explore both factors shaping GC morphology as well as factors dynamically regulated during the late stomatal lineage, we selected 23 targets fitting into the following categories (**Table S4**): (a) stomatal/signaling (6 genes including putative receptors and genes known to affect stomatal patterning), (b) transcription factors (5 predicted or known TFs), (c) cell division/cytoskeleton (7 genes with predicted functions that might affect cell division or the cytoskeleton) and (d) expression dynamics (5 genes selected based on their expression dynamics). For the latter category, we selected 2 genes (GRPL1 and AT5G43450) that normally experience up-regulation during stomatal maturation while 3 genes (i.e., SAUR71, LTL1 and PDF1) normally experience down-regulation during stomatal maturation.

### Electron microscopy

Cotyledons from 12-day-old seedlings were dissected and immediately fixed in 2.5% Glutaraldehyde/ 2% Formaldehyde for 2 h at room temperature. Samples were post-fixed for 2 h in 1% osmium tetroxide (in water) on ice and 1 h in 1% uranyl acetate at room temperature in the dark. Between each incubation step, samples were washed several times with water. After fixation, cotyledons were dehydrated in a graded ethanol series (75%, 90%) followed by 2x acetone (100%), each for 1h. Dehydrated samples were infiltrated with increasing epoxy resin concentrations (10% for 3 h; 25% for 3 h; 50% for 7 h; 75% overnight; 100% for 7 h and 100% overnight). After resin infiltration, samples were embedded in flat embedding molds and polymerized at 60°C for 2 days. Ultrathin sections (50 nm) were performed at the Leica UC7 ultramicrotome. Sections were contrasted with uranyl acetate (in 50% ethanol) and lead citrate. Analysis of stomata ultrastructure was performed at the Jeol JEM-1400Plus transmission electron microscope operated at 120kV and equipped with a 4K CMOS camera TemCam-F416 (Tietz).

### Statistical analysis

Sample sizes for each experiment are specified in the figure legends. For microscopy data, each replicate is a cotyledon or leaf from an independent individual, except for in the opening and closing assays, where each replicate is a stomatal complex. For RNA-seq data, each replicate is a sample containing ∼3,000 cells. Data were analyzed with the statistical software R. For sample comparisons, statistical tests used are indicated in the figure legends.

## Acknowledgements

We would like to thank Rebecca Stahl and Lorenz Henneberg at the ZMBP Microscopy Facility, and the staff at the Stanford Shared FACS Facility for their support.

## Data and resource availability

The raw data for the RNA-seq datasets generated are being made available through Gene Expression Omnibus (GEO) and have been assigned Bioproject number PRJNA1356271.

## Supplementary Figures

**Figure S1.**
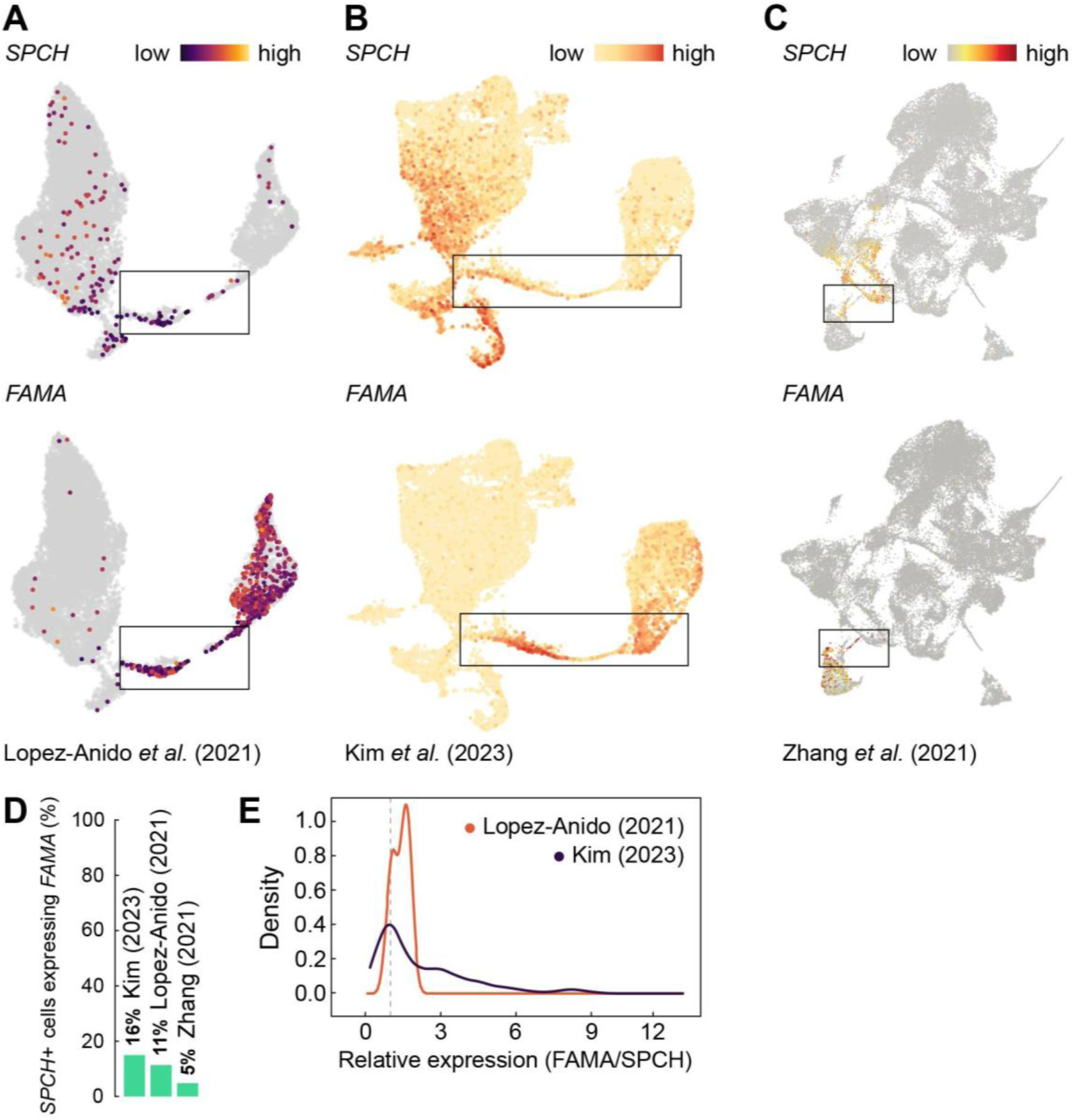
Cells throughout the stomatal lineage co-express *SPCH* and *FAMA*. **A-C.** UMAP plots showing expression of *SPCH* and *FAMA* in three representative scRNA-seq studies: Lopez-Anido *et al*. (2021) (**A**), Kim *et al*. (2023) (**B**) and Zhang *et al*. (2021) (**C**). Boxes highlight late GMCs/young GCs. **D.** Barplot indicating the percentage of *SPCH+* cells that also express *FAMA* across the three different studies. **E.** Density plot of the relative expression between *FAMA* and *SPCH* in cells co-expressing both genes across two studies. The dashed line indicates equal read counts.

**Figure S2.**
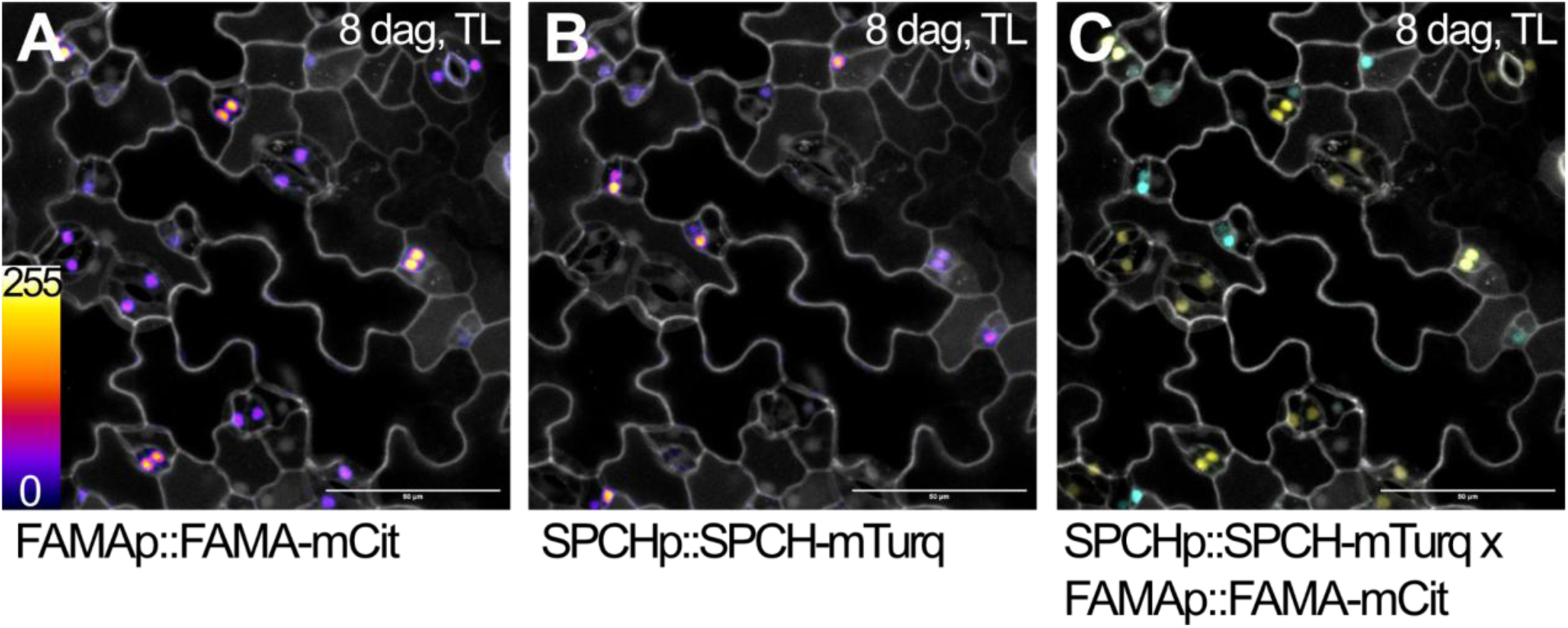
Expression overlap of *SPCH* and *FAMA*. Confocal images showing overlap in expression and protein accumulation between *FAMAp::FAMA-mCit* (**A**), *SPCHp::SPCH-mTurq* (**B**), and merged (**C**) in true leaves of 8 dag seedlings. Membranes are visualized using propidium iodide (white). Scale bars indicate 50 µm.

**Figure S3.**
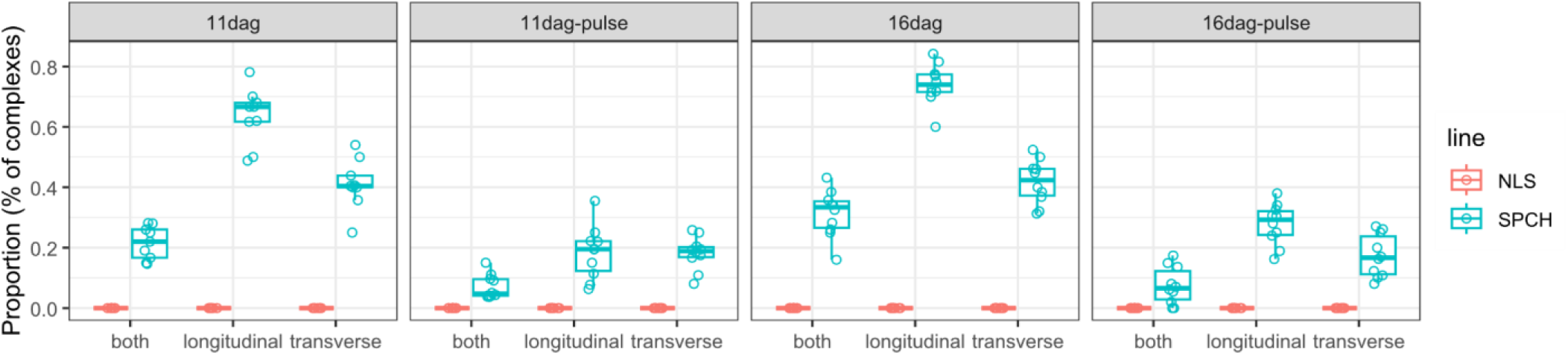
Short ectopic induction of SPCH in the late lineage triggers abnormal guard cell divisions. Quantification of additional divisions upon short FAMAp>>SPCH induction in 11 or 16 days after germination (dag) cotyledons (N=8 leaves). Pulses are treatments with DEX 30 μM for 24h on the 4th day, after which seedlings were returned to normal growth media. Complexes can show additional longitudinal (long.), transverse (trans.) divisions, or both.

**Figure S4.**
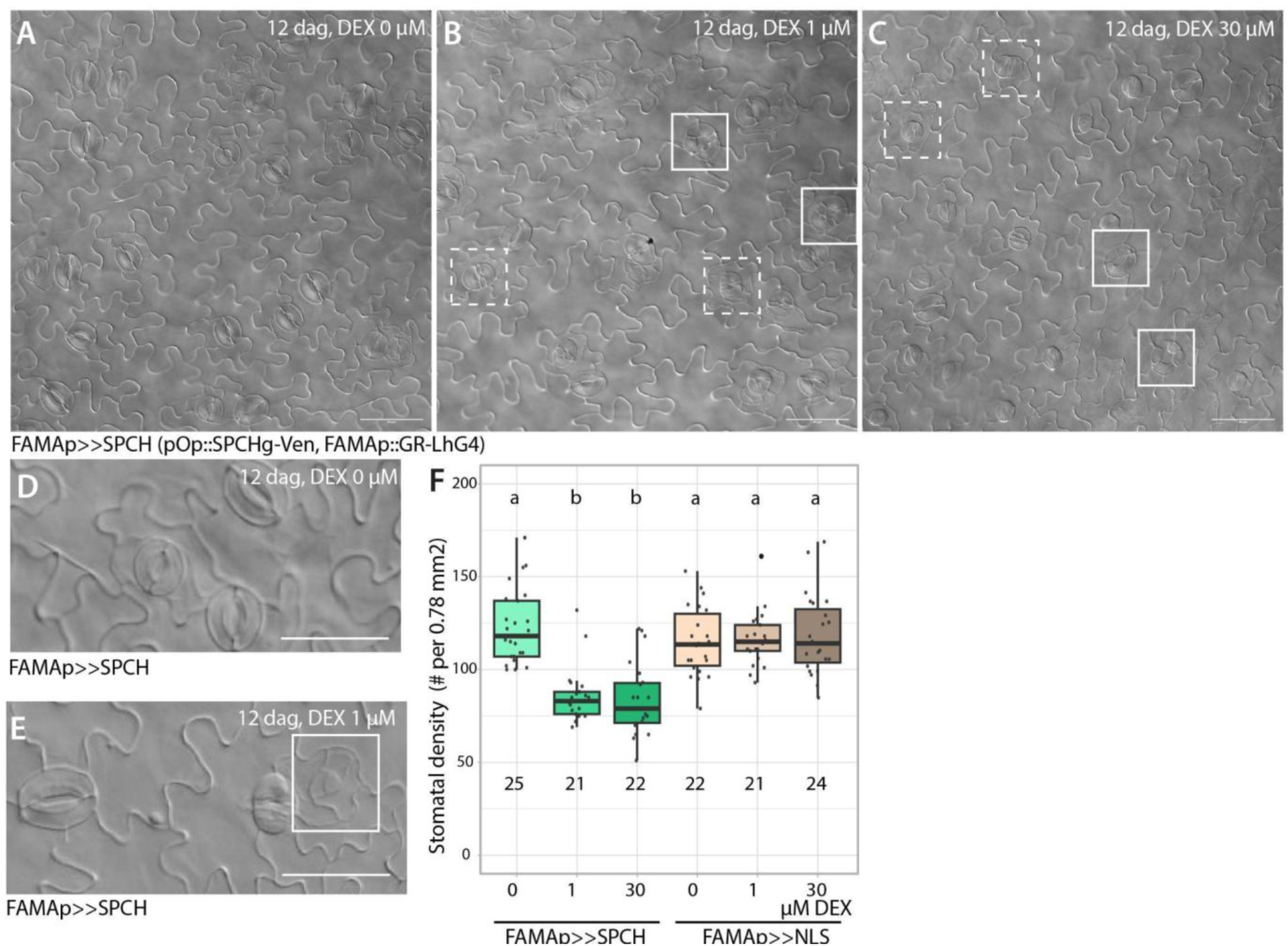
Stomatal phenotypes found in *FAMAp>>SPCH* misexpression lines A-C. DIC images of 12 dag *FAMAp>>SPCH* cotyledons grown without (**A**), with 1 µM (**B**) or 30 µM (**C**) DEX. Abnormal transverse (solid squares) and longitudinal (dashed squares) cell divisions are highlighted. Scale bars indicate 50 µm. **D-E.** Close-ups of aborted stomatal complexes upon *FAMAp>>SPCH* induction. Scale bars indicate 25 µm. **F**. Stomatal density in 12 dag *FAMAp>>SPCH* cotyledons grown in increasingly higher DEX concentration (N=21-25 leaves per sample). Different letters indicate statistical differences (ANOVA, Tukey HSD test, p<0.05).

**Figure S5.**
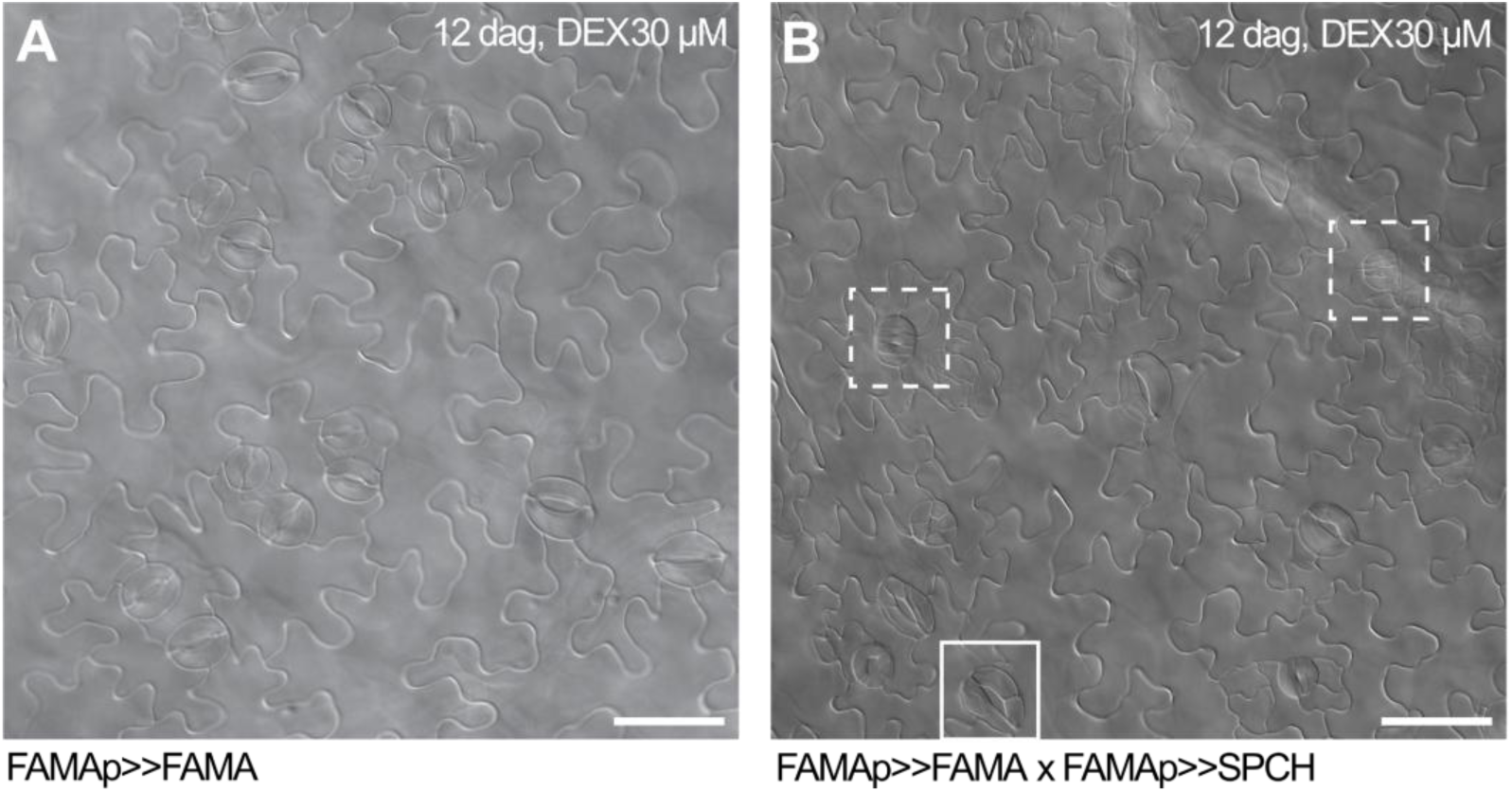
FAMA induction does not abolish SPCH induction phenotypes. DIC images of 12 dag cotyledons upon induction of *FAMAp>>FAMA* (**A**) or combined induction of *FAMAp>>FAMA* and *FAMAp>>SPCH* (**B**). Abnormal transverse (solid squares) and longitudinal (dashed squares) cell divisions are highlighted. Scale bars indicate 50 µm.

**Figure S6.**
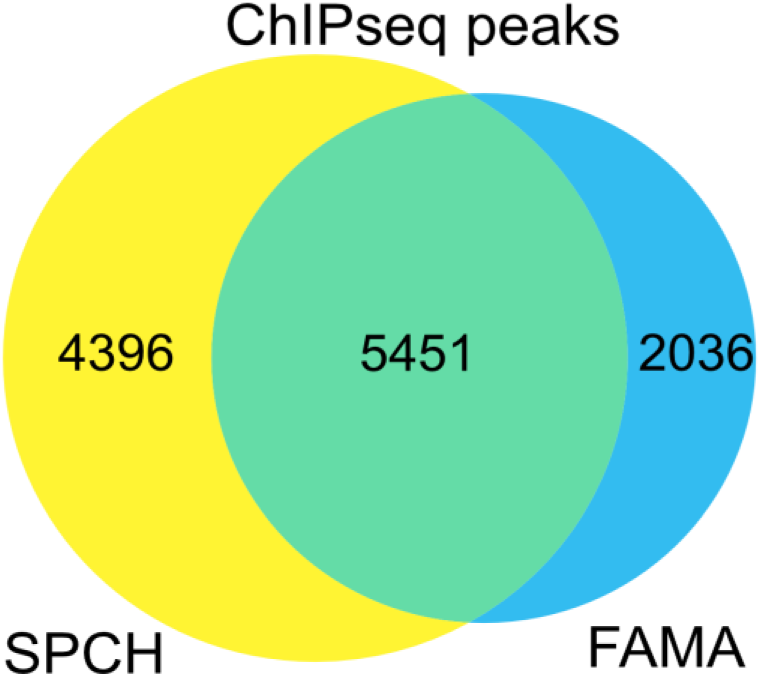
SPCH- and FAMA-bound genes largely overlap. Venn diagram showing the overlap in genes bound by SPCH and FAMA as detected by ChIP-seq, adapted from Liu *et al*., 2024.

**Figure S7.**
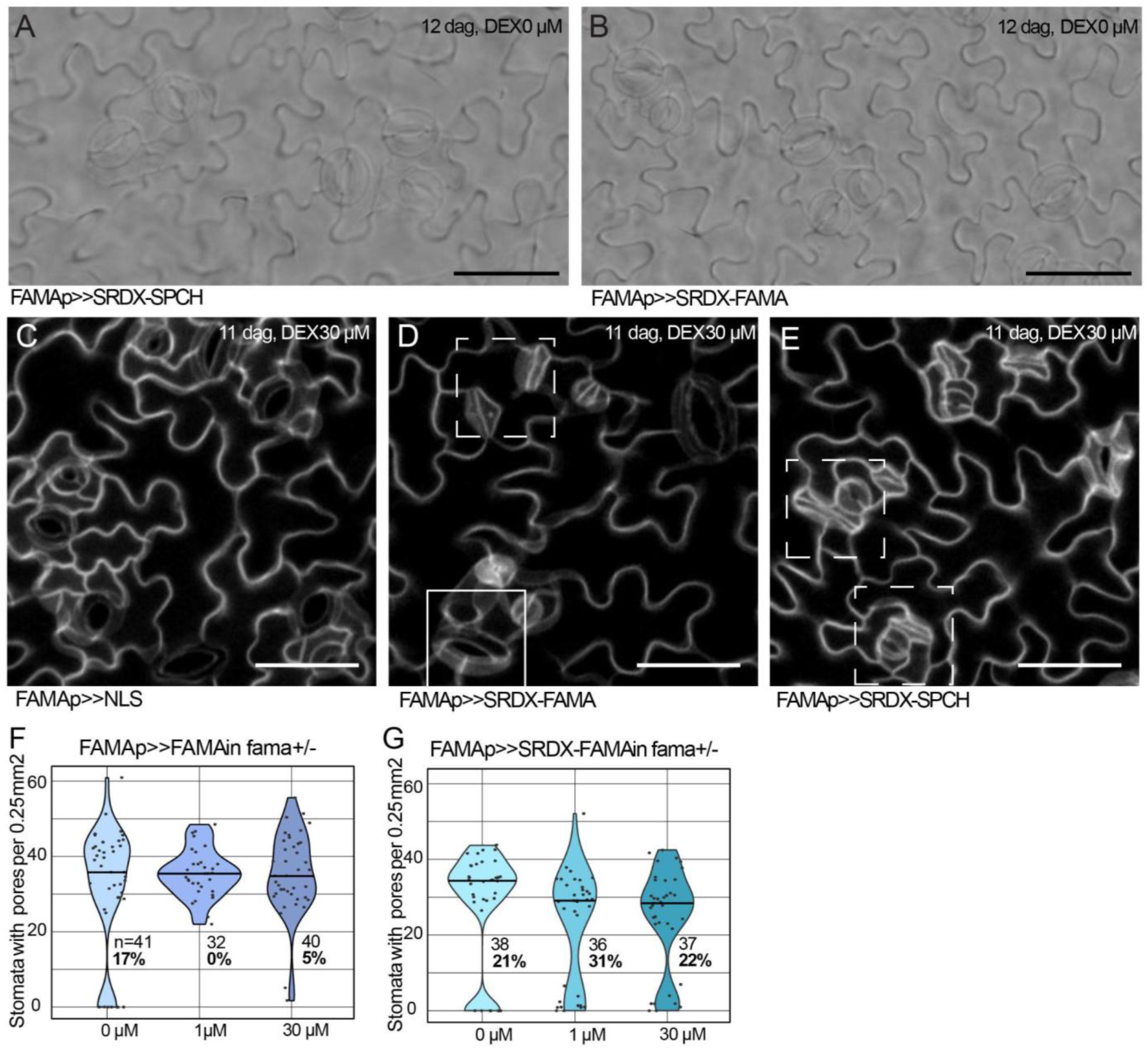
Induction of *FAMA* or *SRDX-FAMA* can rescue the division phenotype of the *fama+/-* mutant. **A-B.** *FAMAp>>SRDX-SPCH* and *FAMAp>>SRDX-FAMA* lines show no changes in stomatal morphology when grown without DEX. DIC images of 12 dag cotyledons, scale bars indicate 50 µm. **C-E.** Confocal stacks of *FAMAp>>NLS, FAMAp>>SRDX-FAMA* and *FAMAp>>SRDX-SPCH* induction lines showing stomatal complexes with additional divisions (solid square) or pavement cell-like morphology (dashed squares) upon SRDX-SPCH or SRDX-FAMA induction. 11 dag cotyledons, scale bars indicate 50 µm. Membranes are visualized using propidium iodide and the plasma membrane marker *ML1p::mCherry-RCl2A* (white). **F.** Number of stomata with pores in *fama+/-*grown without or with 1 µM or 30 µM DEX inducing expression of *FAMAp>>FAMA*. N=32-41 leaves. % indicates the percentage of leaves that have fewer than 10 pore-containing complexes, expected ∼25% in *fama+/-*. **G.** Number of stomata with pores in *fama+/-* grown in half-strength MS media or with 1 µM or 30 µM DEX inducing expression of *FAMAp>>SRDX-FAMA*. N=36-38 leaves. % indicates the percentage of leaves that have fewer than 10 pore-containing complexes, expected ∼25% in *fama+/-*.

**Figure S8.**
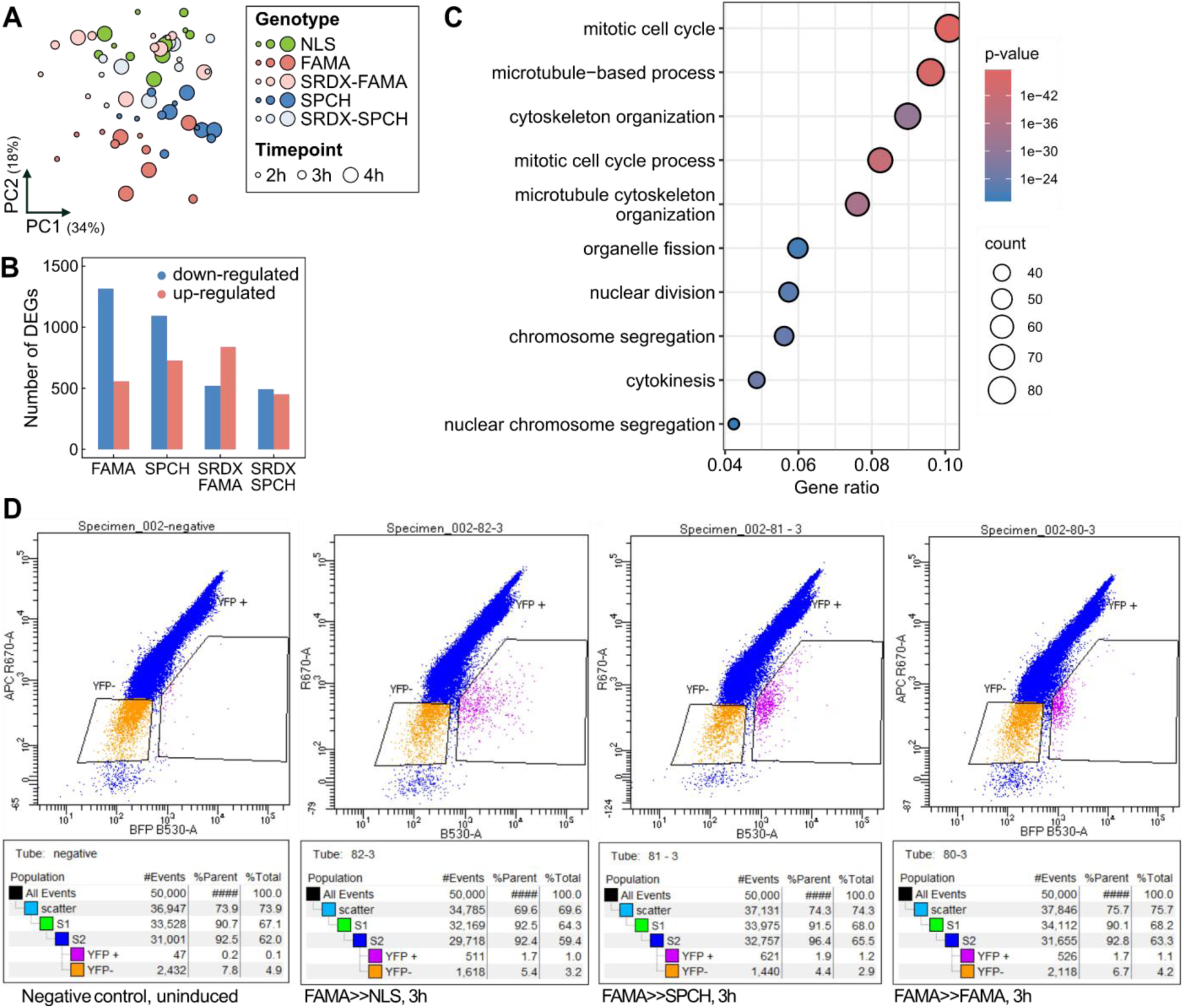
RNA-seq of *FAMA+* cells shortly after NLS and transgene induction reveals high variability and differing cell composition. **A.** Principal component analysis (PCA) plot of mRNA-sequenced samples. Dot size indicates the timepoint and colour the genotype. **B.** Barplot showing the number of genes differentially expressed upon induction of *FAMA*, *SPCH*, *SRDX-FAMA* or *SRDX-SPCH* compared to control *NLS* induction. **C.** Gene ontology (GO) enrichment analysis in genes down-regulated upon *SPCH* induction compared to *NLS* induction highlighting unexpected GO terms related to cell division. **D.** Representative flow cytometry plots from sorting Venus positive cells. Red and yellow fluorescence channels on y-axis and x-axis respectively. Comparing an uninduced sample as control (left) and three induction lines after 3 hours of DEX incubation shows increased yellow fluorescence resulting in cells shifting to the right. The right window was used for collection.

**Figure S9.**
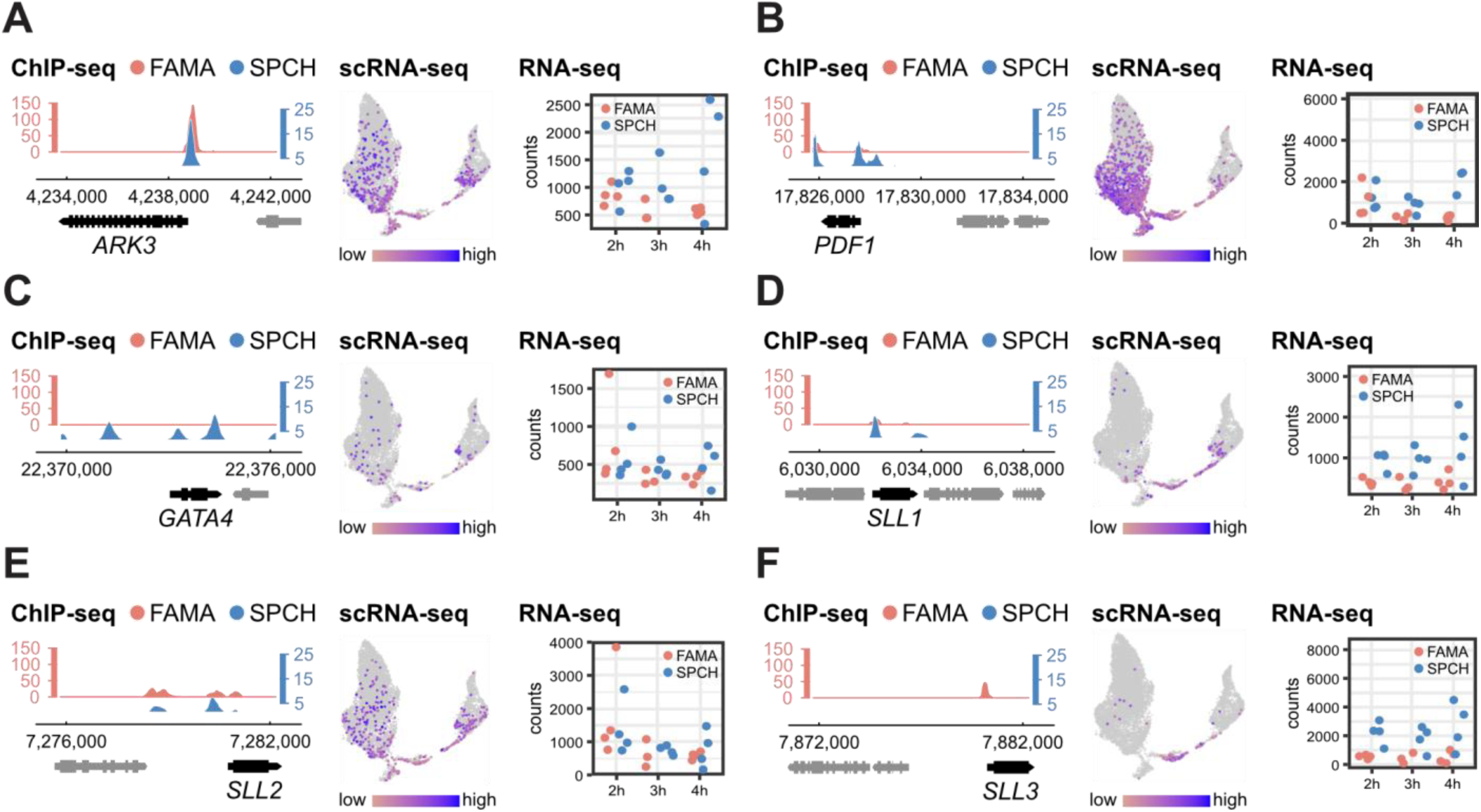
Available information from omics studies about additional candidates. Information used for our target gene selection for other key genes mentioned throughout this manuscript: *ARK3* (**A**), *PDF1* (**B**), *GATA4* (**C**), *SLL1* (**D**), *SLL2* (**E**) and *SLL3* (**F**). ChIP-seq peaks for SPCH and FAMA are on the left (Lau et al., 2014; Liu et al., 2024). Plots showing expression in stomatal scRNA-seq are in the middle (Lopez-Anido et al., 2021). Expression plots highlighting transcriptional changes upon SPCH and FAMA induction in our study are on the right.

**Figure S10.**
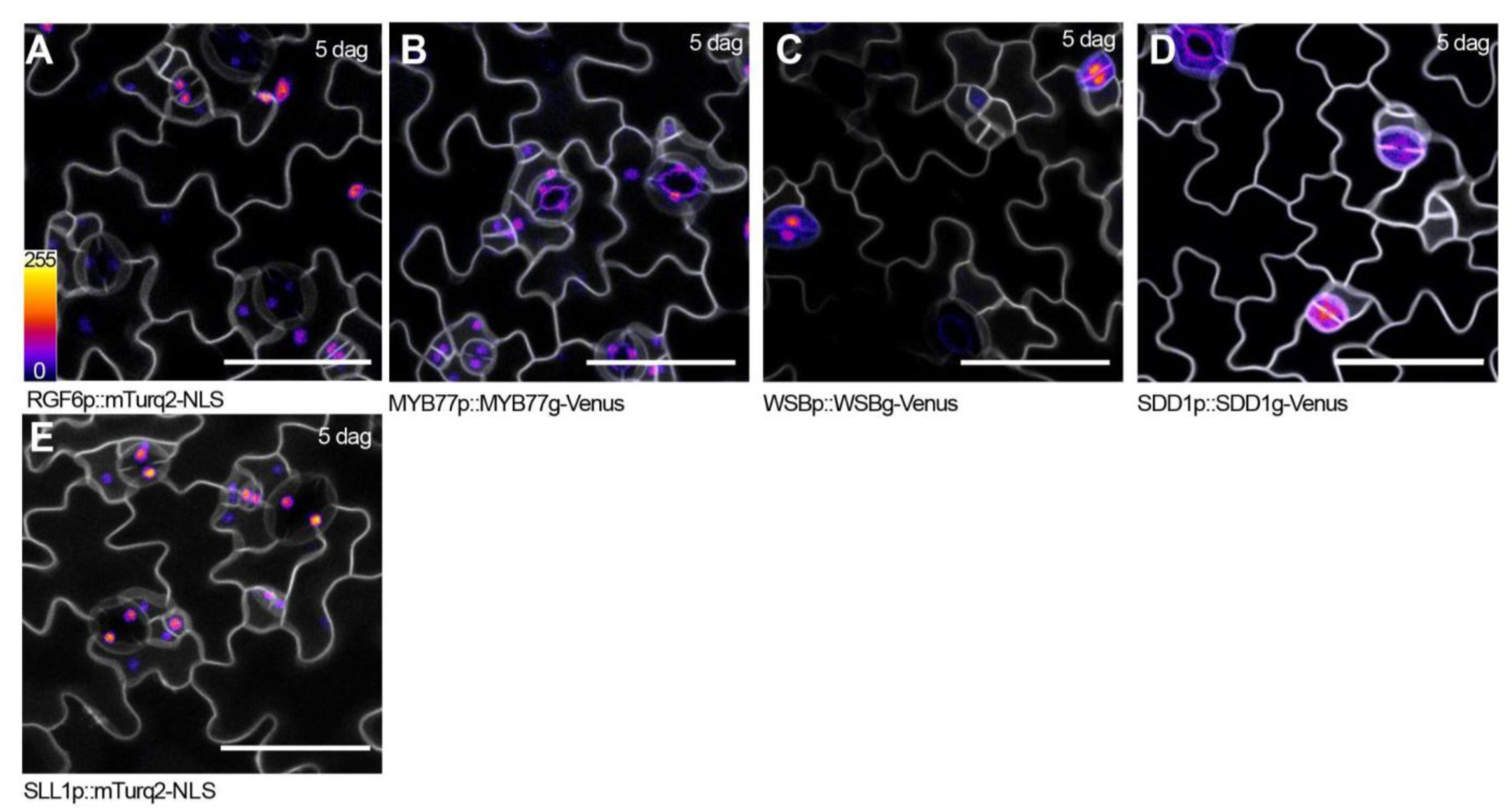
Expression patterns of SPCH/FAMA targets during the late stomatal lineage A-E. Confocal images of transcriptional and translational reporters for putative targets of SPCH and FAMA during the late lineage: *RGF6* (**A**), *MYB77* (**B**), *WSB* (**C**), *SDD1* (**D**) and *SLL1* (**E**) in 5 dag cotyledons. Scale bars indicate 50 µm. Membranes are visualized using the plasma membrane marker *ML1p::mCherry-RCl2A* (white).

**Figure S11.**
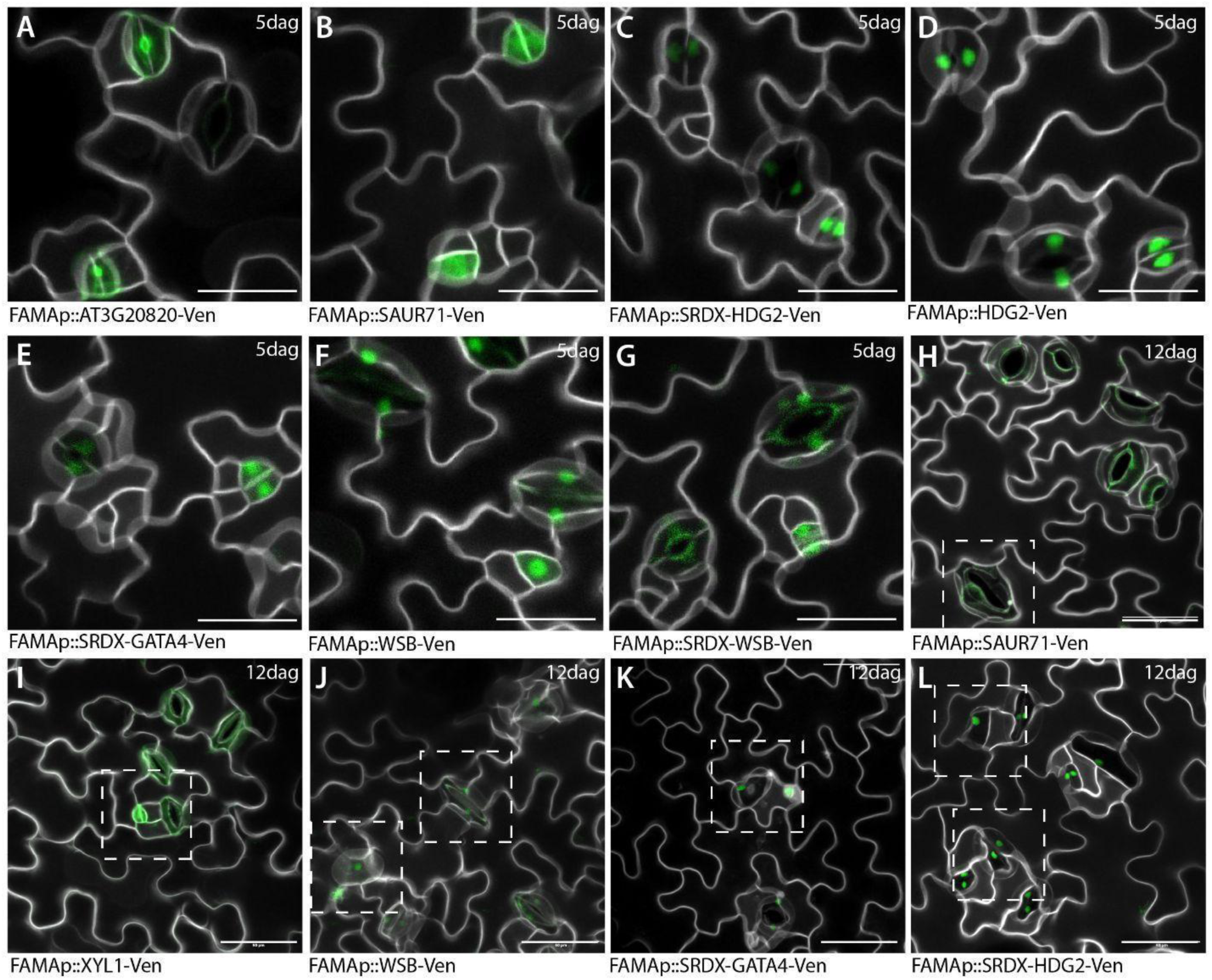
Subcellular localization of misexpressed putative targets of SPCH and FAMA. Confocal images of misexpression lines for selected putative targets of SPCH and FAMA in 5 dag (**A-G**) or 12 dag (**H-L**) cotyledons. Dashed squares indicate abnormal stomatal morphology. Scale bars indicate 50 µm. Membranes are visualized using propidium iodide and the plasma membrane marker *ML1p::mCherry-RCl2A* (white).

**Figure S12.**
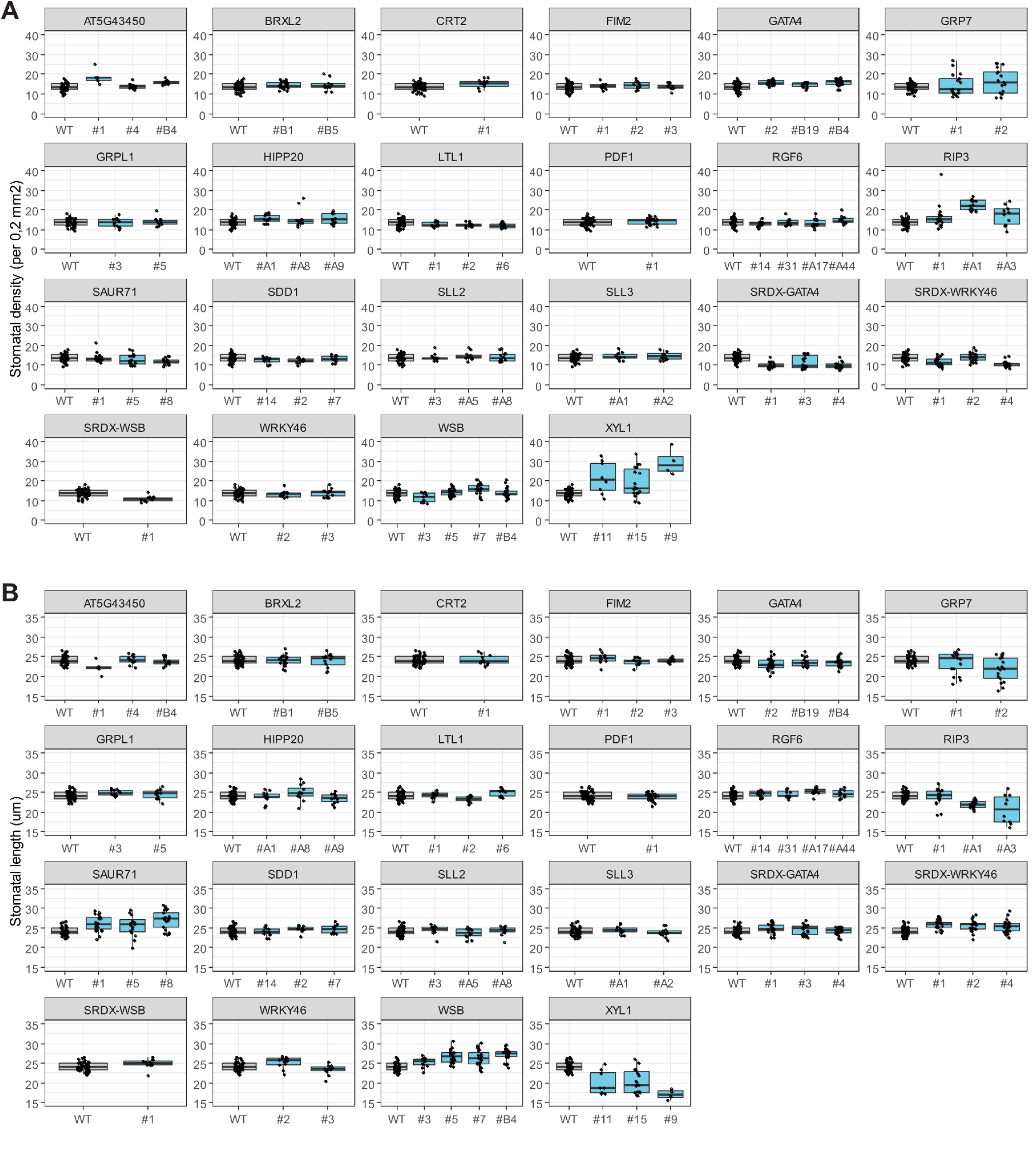
Changes in stomatal density and size in lines misexpressing putative targets of SPCH and FAMA in the late lineage. Stomatal density (**A**) and stomatal length (**B**) in 12 dag cotyledons of lines misexpressing putative targets of SPCH and FAMA. Each boxplot corresponds to an independent T2 line.

**Figure S13.**
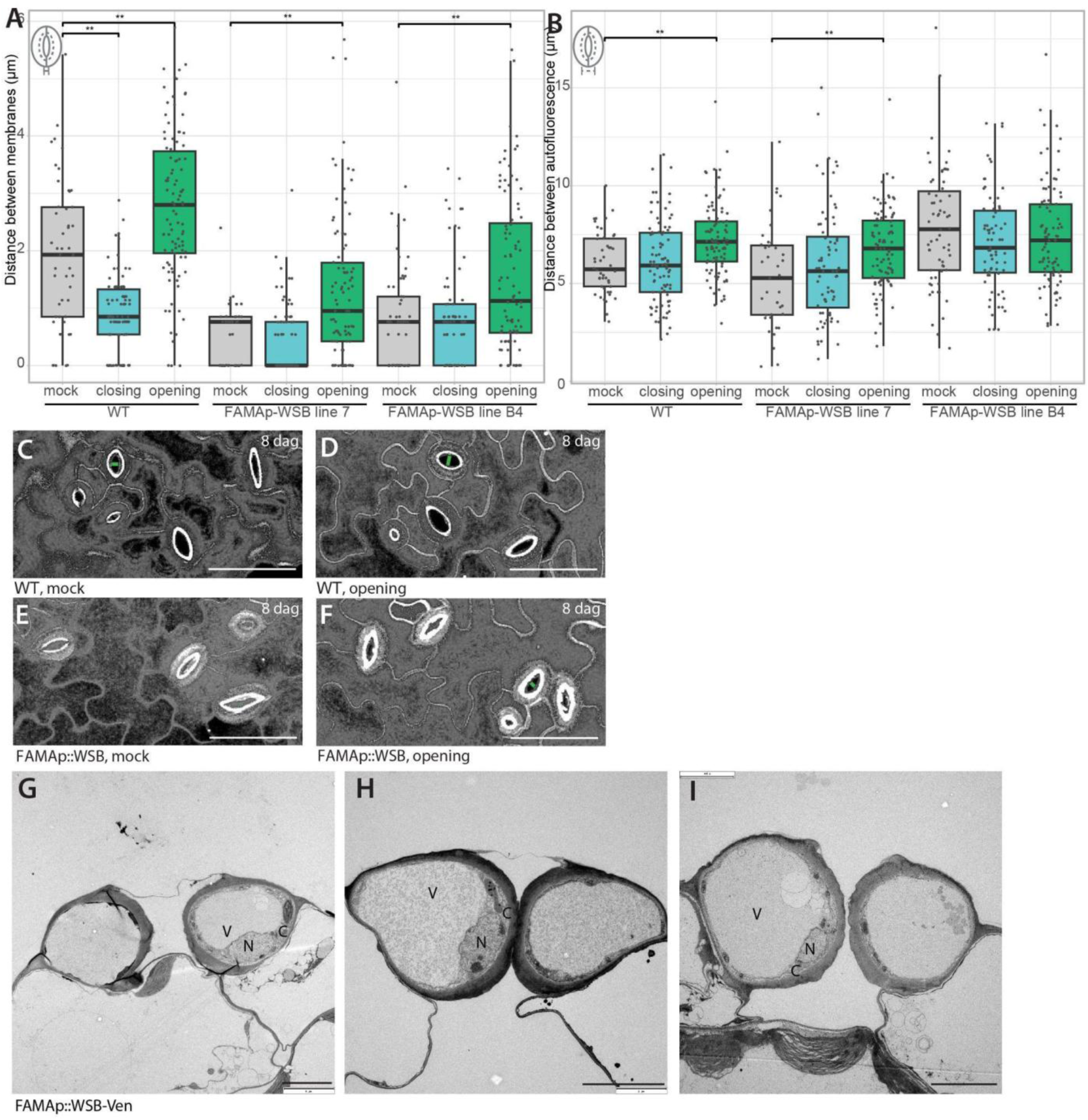
Misexpression of *WSB* impairs stomatal opening and changes in physical cell morphology. **A-B.** Longest distance between guard cell membranes (**A**) and between pore autofluorescences (**B**) in 8 dag cotyledons of wild-type and two independent *FAMAp::WSB* lines exposed to closing solution, opening solutions, or mock. Asterisks indicate statistical differences (Student’s t test, ** p<0.01, *** p<0.005)(N=39-98 stomata across 3-5 leaves). **C-F.** Representative confocal z-stacks of wild-type (**C-D**) and *FAMAp::WSB* (**E-F**) exposed to mock or opening solutions. Membranes are visualized using propidium iodide and the plasma membrane marker *ML1p::mCherry-RCl2A* (white). Black areas indicate that no membrane was present throughout the stack. Green lines indicate where distance between guard cell membranes was measured. Scale bars indicate 50 µm. **G-I.** Cross-sectional TEM images showing large guard cells in *FAMAp::WSB*. V =vacuole , N =nucleus , C =cytosol , Scale bars indicate 5 µm.

**Figure S14.**
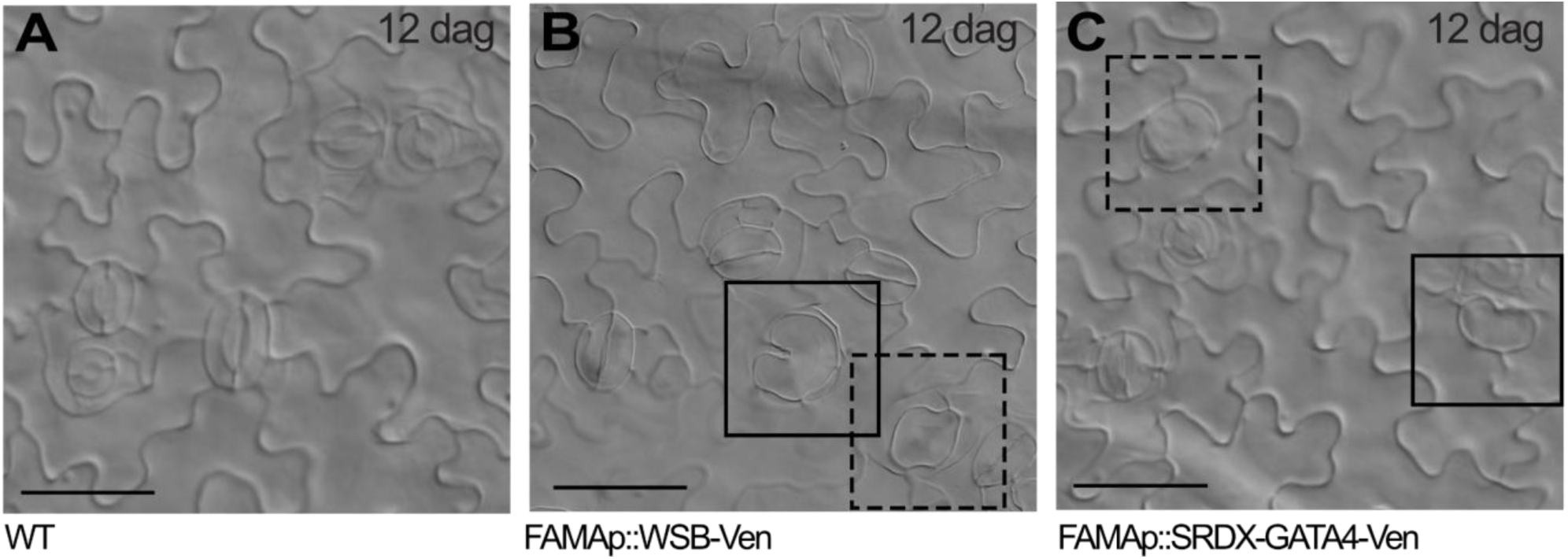
Misexpression of a subset of putative targets of SPCH and FAMA triggers the formation of single guard cells. DIC images of 12 dag cotyledons of wild-type (**A**), *FAMAp::WSB-Ven* (**B**) and *FAMAp::SRDX-GATA4-Ven* (**C**). Solid squares highlight large kidney-shaped single GCs and dashed squares indicate large round single GCs. Scale bars indicate 50 µm.

**Figure S15.**
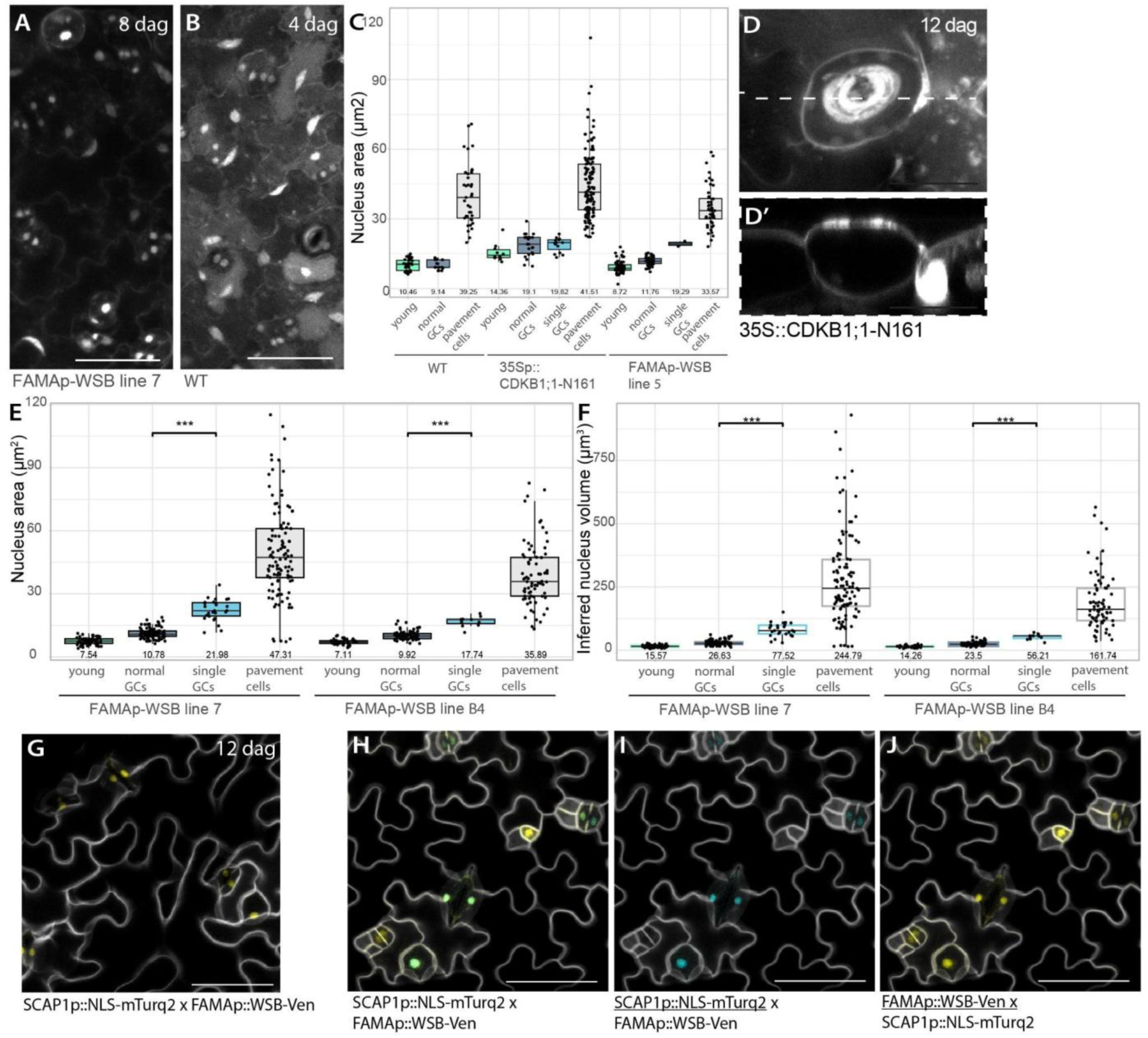
Single guard cells induced by *WSB* misexpression differentiate after DNA replication. **A-B.** Confocal z-stacks of Hoechst stained *FAMAp::WSB* and WT leaves used for nucleus area measurements. Scale bars indicate 50 µm. **C.** Graph showing nucleus area of young stomatal cells, regular GCs, SGCs (single guard cells), and pavement cells at 4 dag for WT, *35S::CDKB1;1-N161*, and *FAMAp::WSB*. **D.** Confocal image of large single cells in 12 dag cotyledons of *CDKB1;1-N161*. Scale bars indicate 25 µm. **E.** Additional nucleus area measurements for 2 independent *FAMAp::WSB* lines, expanding in Figure 5E. **F.** Nucleus volume inferred from area measurements in E. **G.** *FAMAp::WSB* signal for the corresponding image in Figure 5D. **H-I**. Additional images showing SCAP reporter expression in *FAMAp::WSB.* Scale bars indicate 50 µm.

