## Supplemental Figures for "Targets of SPEECHLESS and FAMA control guard cell division and expansion in the late stomatal lineage"

### Supplementary Figures and Tables

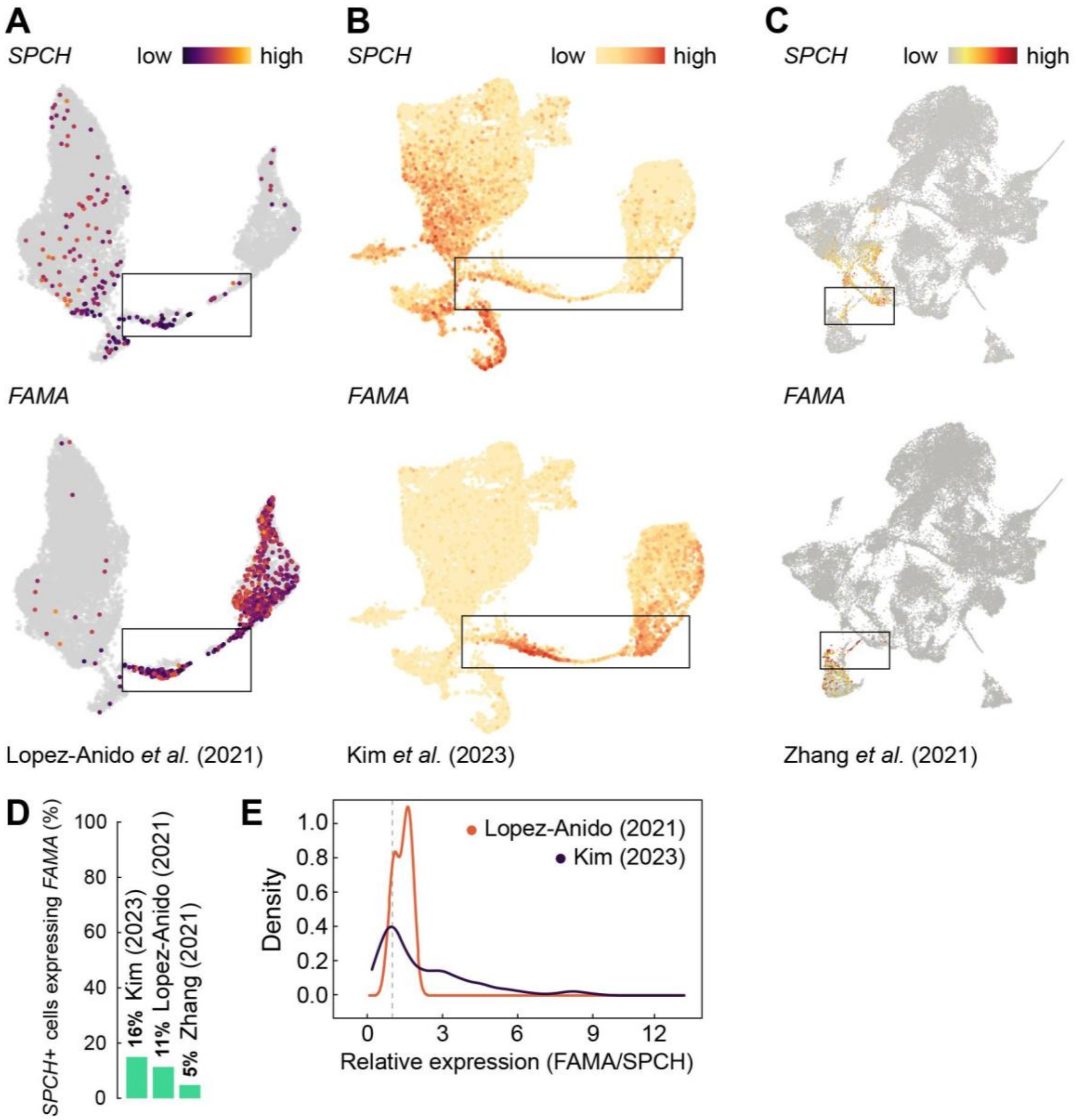


**Figure S1. Cells throughout the stomatal lineage co-express *SPCH* and *FAMA***
**A-C.** UMAP plots showing expression of *SPCH* and *FAMA* in three representative scRNA-seq studies: Lopez-Anido *et al.* (2021) (**A**), Kim *et al.* (2023) (**B**) and Zhang *et al.* (2021) (**C**). Boxes highlight late GMCs/young GCs. **D.** Barplot indicating the percentage of *SPCH+* cells that also express *FAMA* across the three different studies. **E.** Density plot of the relative expression between *FAMA* and *SPCH* in cells co-expressing both genes across two studies. The dashed line indicates equal read counts.

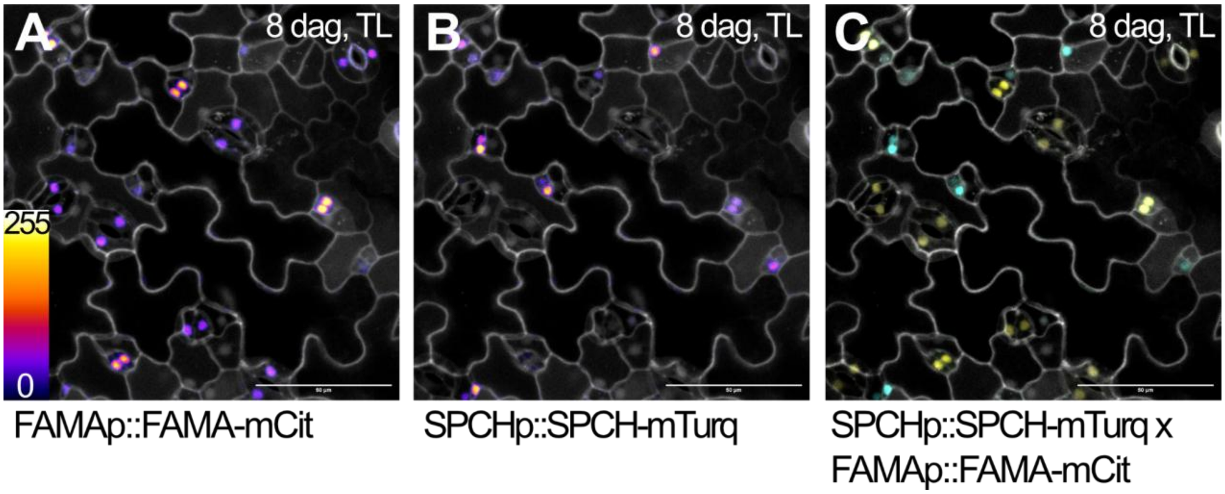


**Figure S2. Expression overlap of *SPCH* and *FAMA***
Confocal images showing overlap in expression and protein accumulation between *FAMAp::FAMA-mCit* (**A**), *SPCHp::SPCH-mTurq* (**B**), and merged (**C**) in true leaves of 8 dag seedlings. Membranes are visualized using propidium iodide (white). Scale bars indicate 50 μm.

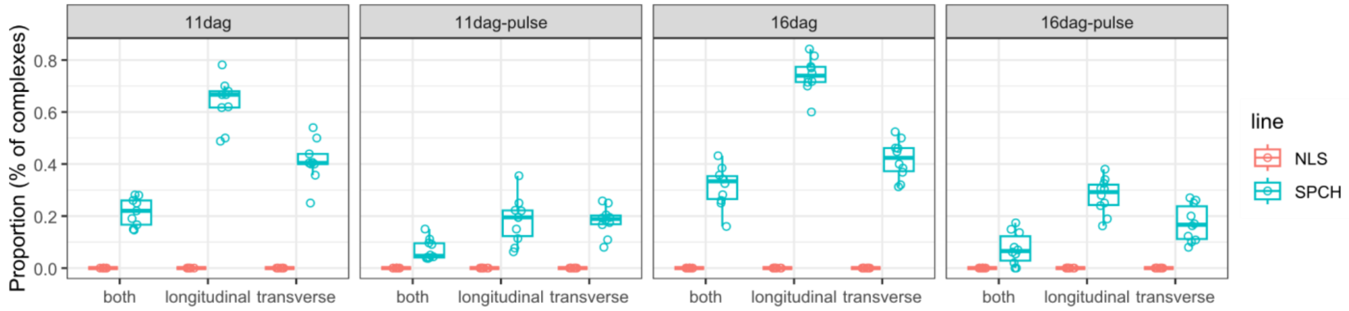


**Figure S3. Short ectopic induction of SPCH in the late lineage triggers abnormal guard cell divisions**

Quantification of additional divisions upon short FAMAp>>SPCH induction in 11 or 16 days after germination (dag) cotyledons (N=8 leaves). Pulses are treatments with DEX 30 μM for 24h on the 4th day, after which seedlings were returned to normal growth media. Complexes can show additional longitudinal (long.), transverse (trans.) divisions, or both.

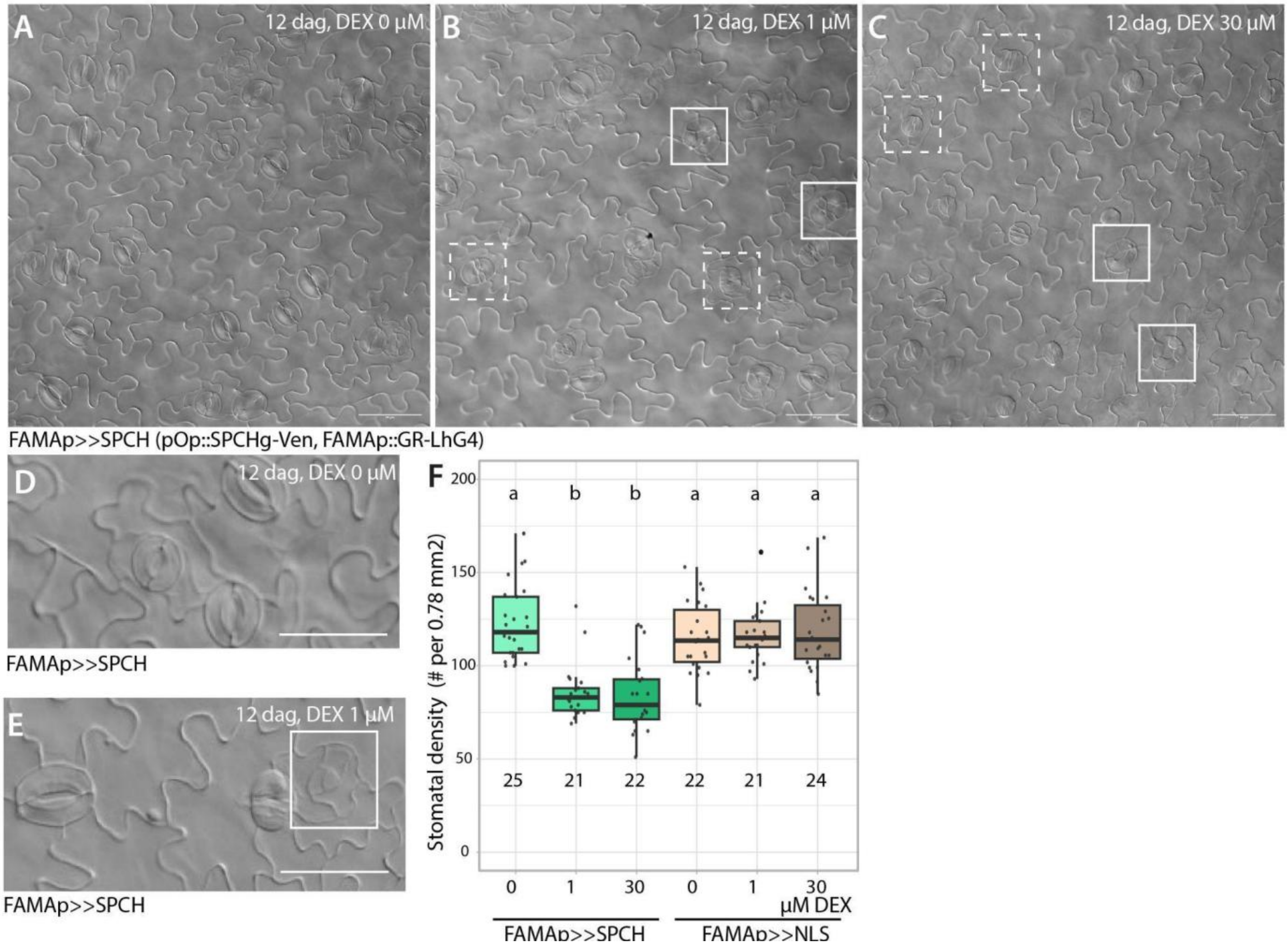


**Figure S4. Stomatal phenotypes found in *FAMAp>>SPCH* misexpression lines**
**A-C.** DIC images of 12 dag *FAMAp>>SPCH* cotyledons grown without (**A**), with 1 µM (**B**) or 30 µM (**C**) DEX. Abnormal transverse (solid squares) and longitudinal (dashed squares) cell divisions are highlighted. Scale bars indicate 50 µm. **D-E.** Close-ups of aborted stomatal complexes upon *FAMAp>>SPCH* induction. Scale bars indicate 25 µm. **F**. Stomatal density in 12 dag *FAMAp>>SPCH* cotyledons grown in increasingly higher DEX concentration (N=21-25 leaves per sample). Different letters indicate statistical differences (ANOVA, Tukey HSD test, $p<0.05$).

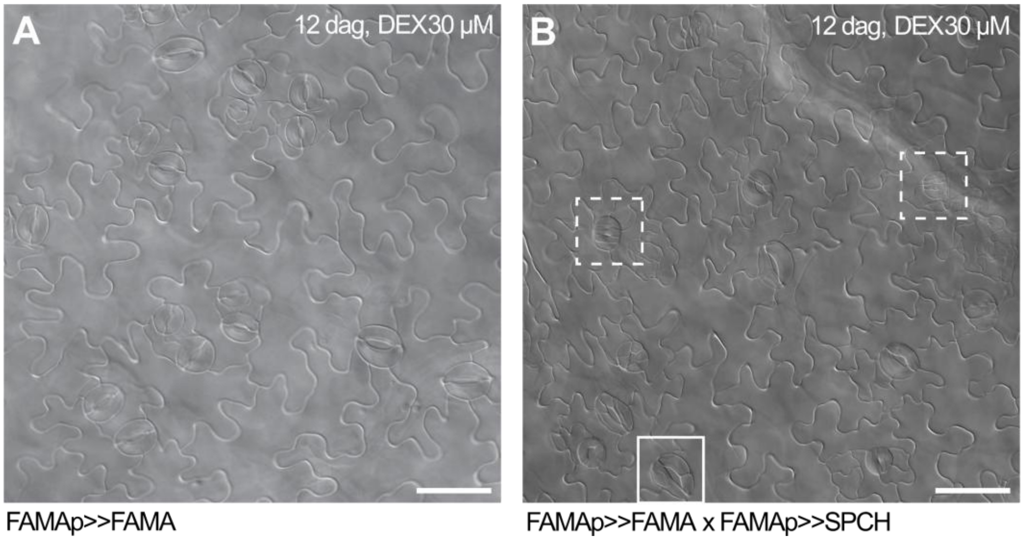


**Figure S5. FAMA induction does not abolish SPCH induction phenotypes**

DIC images of 12 dag cotyledons upon induction of *FAMAp>>FAMA* (**A**) or combined induction of *FAMAp>>FAMA* and *FAMAp>>SPCH* (**B**). Abnormal transverse (solid squares) and longitudinal (dashed squares) cell divisions are highlighted. Scale bars indicate 50 µm.

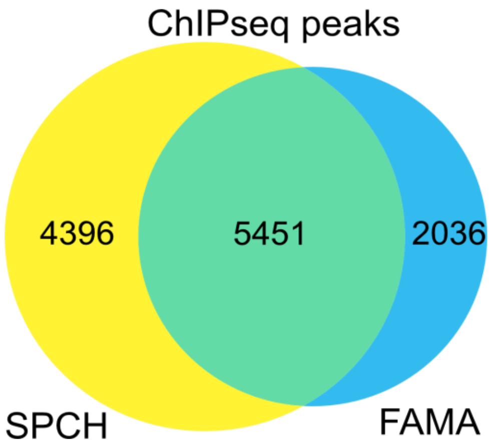


**Figure S6. SPCH- and FAMA-bound genes largely overlap**

Venn diagram showing the overlap in genes bound by SPCH and FAMA as detected by ChIP-seq, adapted from Liu *et al.*, 2024.

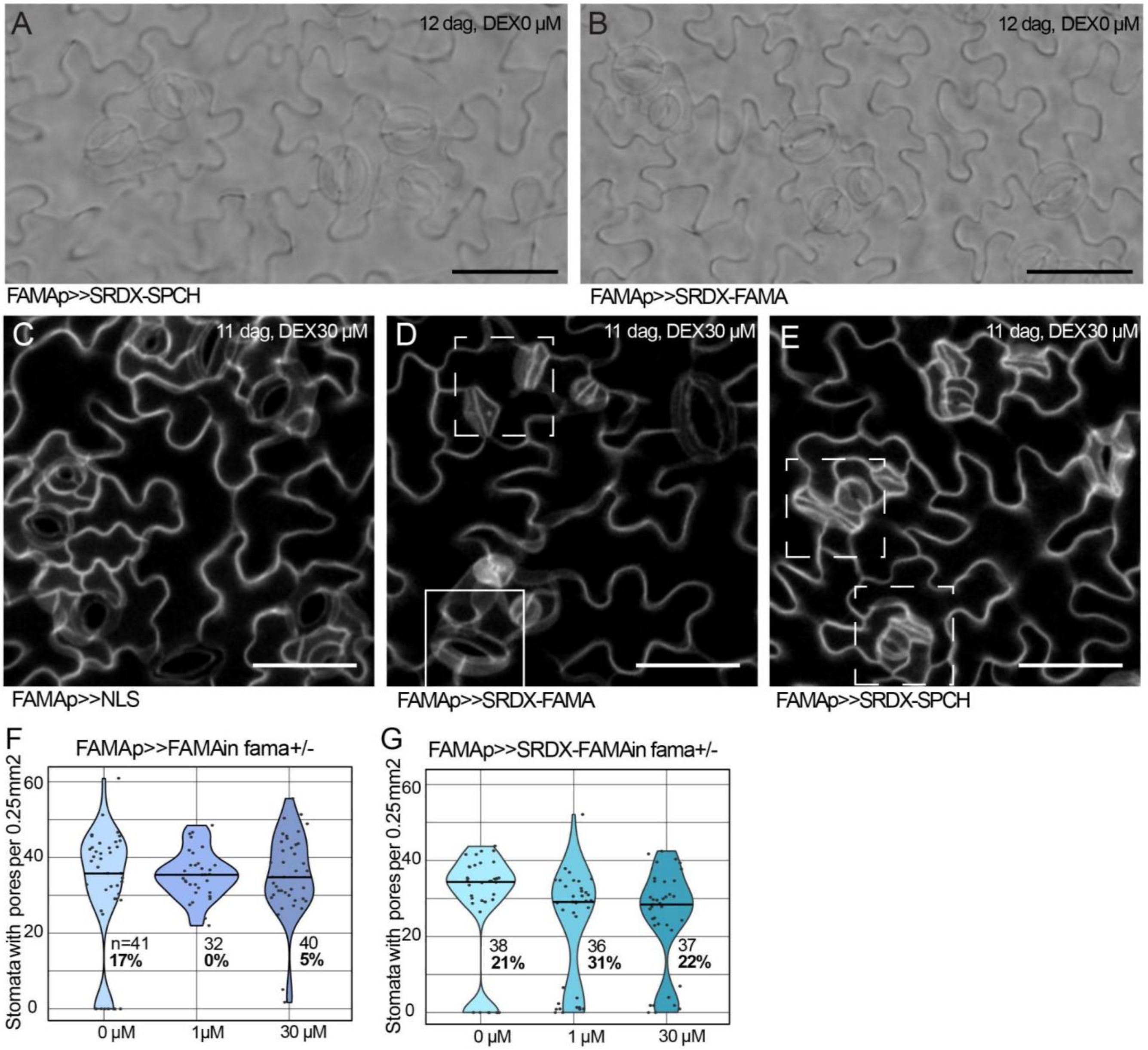


**Figure S7. Induction of *FAMA* or *SRDX-FAMA* can rescue the division phenotype of the *fama+/-* mutant**

**A-B.** *FAMAp>>SRDX-SPCH* and *FAMAp>>SRDX-FAMA* lines show no changes in stomatal morphology when grown without DEX. DIC images of 12 dag cotyledons, scale bars indicate 50 µm. **C-E.** Confocal stacks of *FAMAp>>NLS, FAMAp>>SRDX-FAMA* and *FAMAp>>SRDX-SPCH* induction lines showing stomatal complexes with additional divisions (solid square) or pavement cell-like morphology (dashed squares) upon SRDX-SPCH or SRDX-FAMA induction. 11 dag cotyledons, scale bars indicate 50 μm. Membranes are visualized using propidium iodide and the plasma membrane marker *ML1p::mCherry-RCl2A* (white). **F.** Number of stomata with pores in *fama+/-* grown without or with 1 μM or 30 μM DEX inducing expression of *FAMAp>>FAMA*. N=32-41 leaves. % indicates the percentage of leaves that have fewer than 10 pore-containing complexes, expected ~25% in *fama+/-*. **G.** Number of stomata with pores in *fama+/-* grown in half-strength MS media or with 1 μM or 30 μM DEX inducing expression of *FAMAp>>SRDX-FAMA*. N=36-38 leaves. % indicates the percentage of leaves that have fewer than 10 pore-containing complexes, expected ~25% in *fama+/-*.

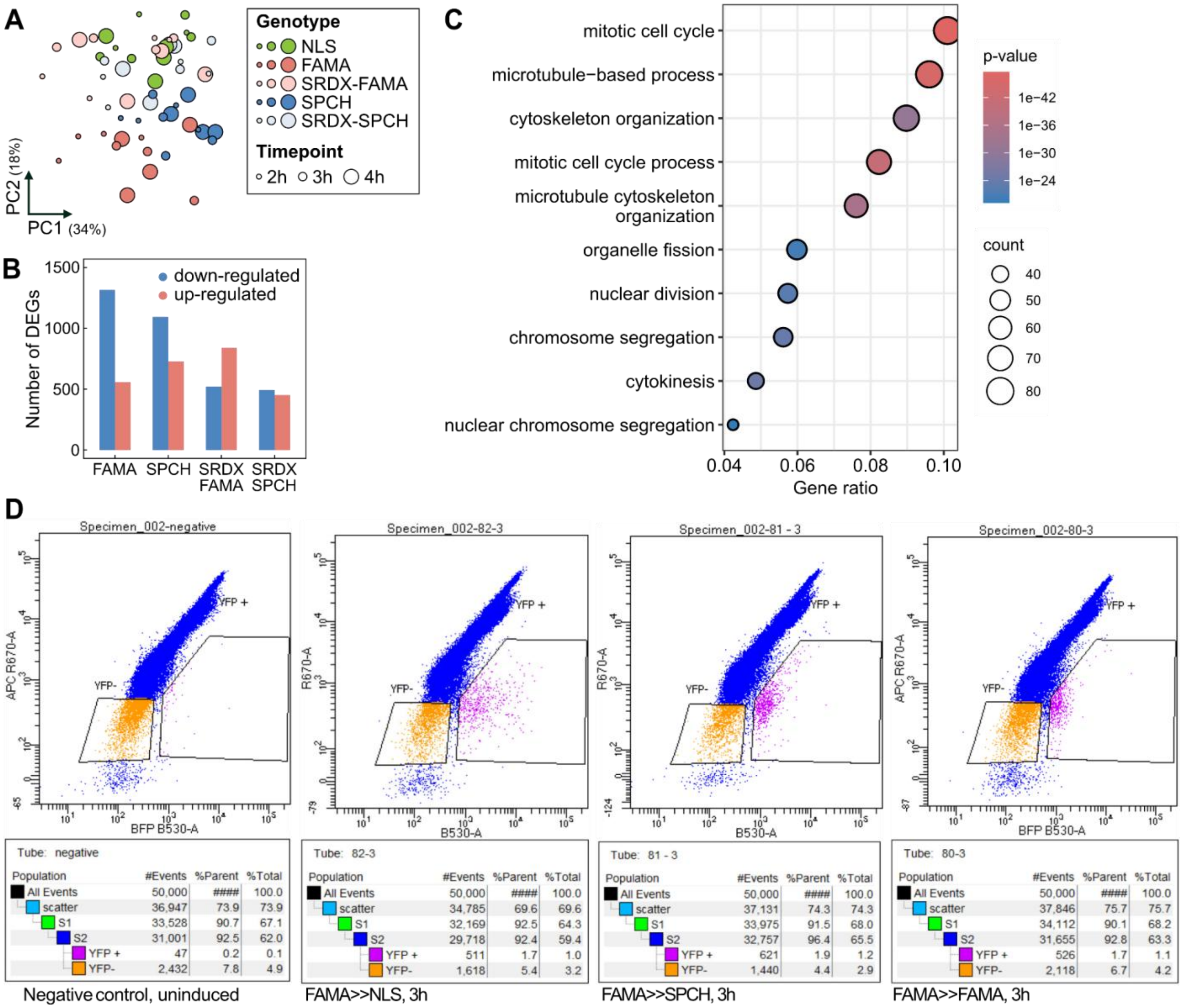


**Figure S8. RNA-seq of *FAMA*+ cells shortly after NLS and transgene induction reveals high variability and differing cell composition**

**A.** Principal component analysis (PCA) plot of mRNA-sequenced samples. Dot size indicates the timepoint and colour the genotype. **B.** Barplot showing the number of genes differentially expressed upon induction of *FAMA*, *SPCH*, *SRDX-FAMA* or *SRDX-SPCH* compared to control *NLS* induction. **C.** Gene ontology (GO) enrichment analysis in genes down-regulated upon *SPCH* induction compared to *NLS* induction highlighting unexpected GO terms related to cell division. **D.** Representative flow cytometry plots from sorting Venus positive cells. Red and yellow fluorescence channels on y-axis and x-axis respectively. Comparing an uninduced sample as control (left) and three induction lines after 3 hours of DEX incubation shows increased yellow fluorescence resulting in cells shifting to the right. The right window was used for collection.

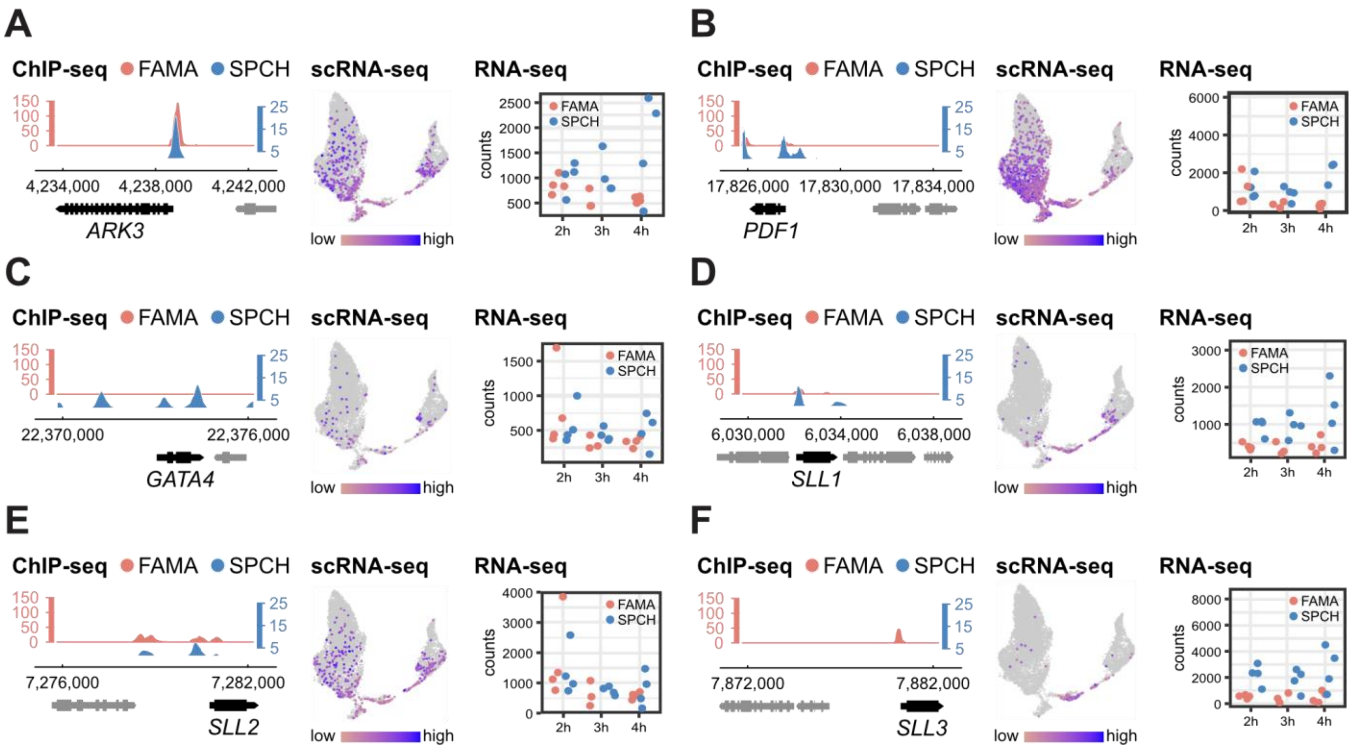


**Figure S9. Available information from omics studies about additional candidates**

Information used for our target gene selection for other key genes mentioned throughout this manuscript: *ARK3* (**A**), *PDF1* (**B**), *GATA4* (**C**), *SLL1* (**D**), *SLL2* (**E**) and *SLL3* (**F**). ChIP-seq peaks for SPCH and FAMA are on the left (Lau et al., 2014; Liu et al., 2024). Plots showing expression in stomatal scRNA-seq are in the middle (Lopez-Anido et al., 2021). Expression plots highlighting transcriptional changes upon SPCH and FAMA induction in our study are on the right.

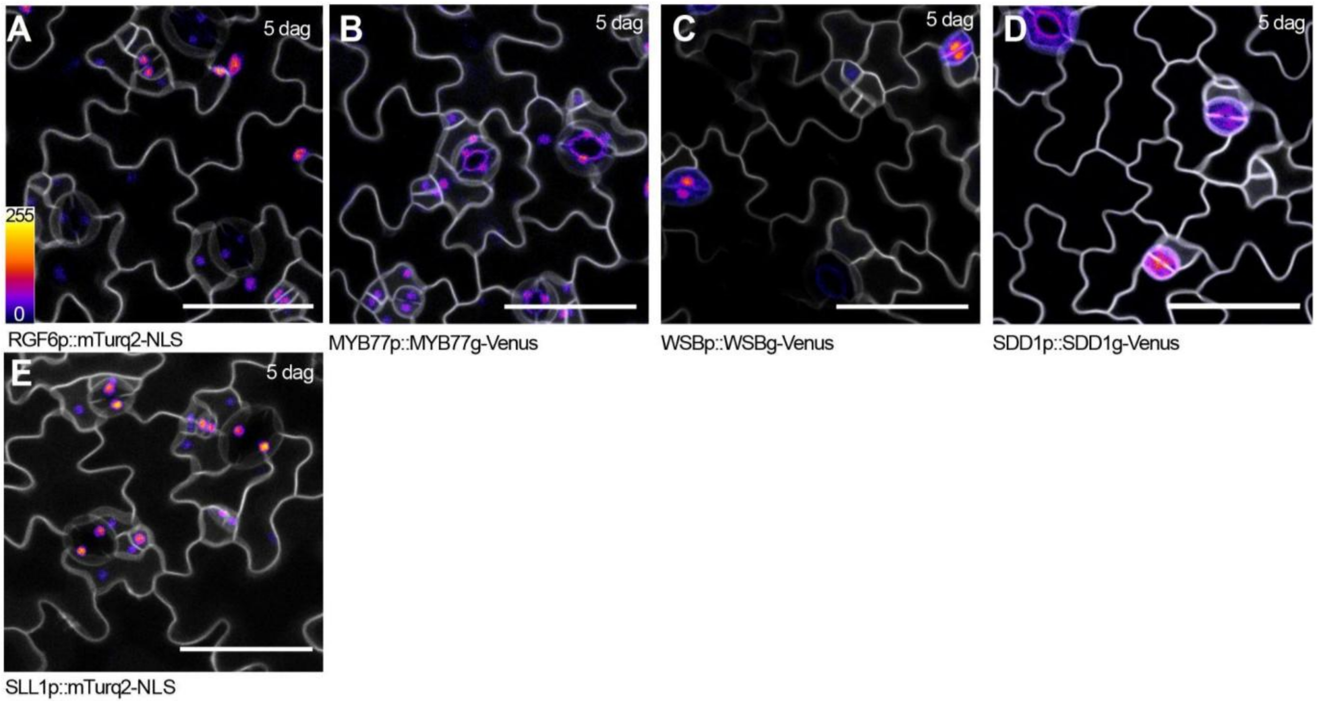


**Figure S10. Expression patterns of SPCH/FAMA targets during the late stomatal lineage**
**A-E.** Confocal images of transcriptional and translational reporters for putative targets of SPCH and FAMA during the late lineage: *RGF6* (**A**), *MYB77* (**B**), *WSB* (**C**), *SDD1* (**D**) and *SLL1* (**E**) in 5 dag cotyledons. Scale bars indicate 50 µm. Membranes are visualized using the plasma membrane marker *ML1p::mCherry-RCl2A* (white).

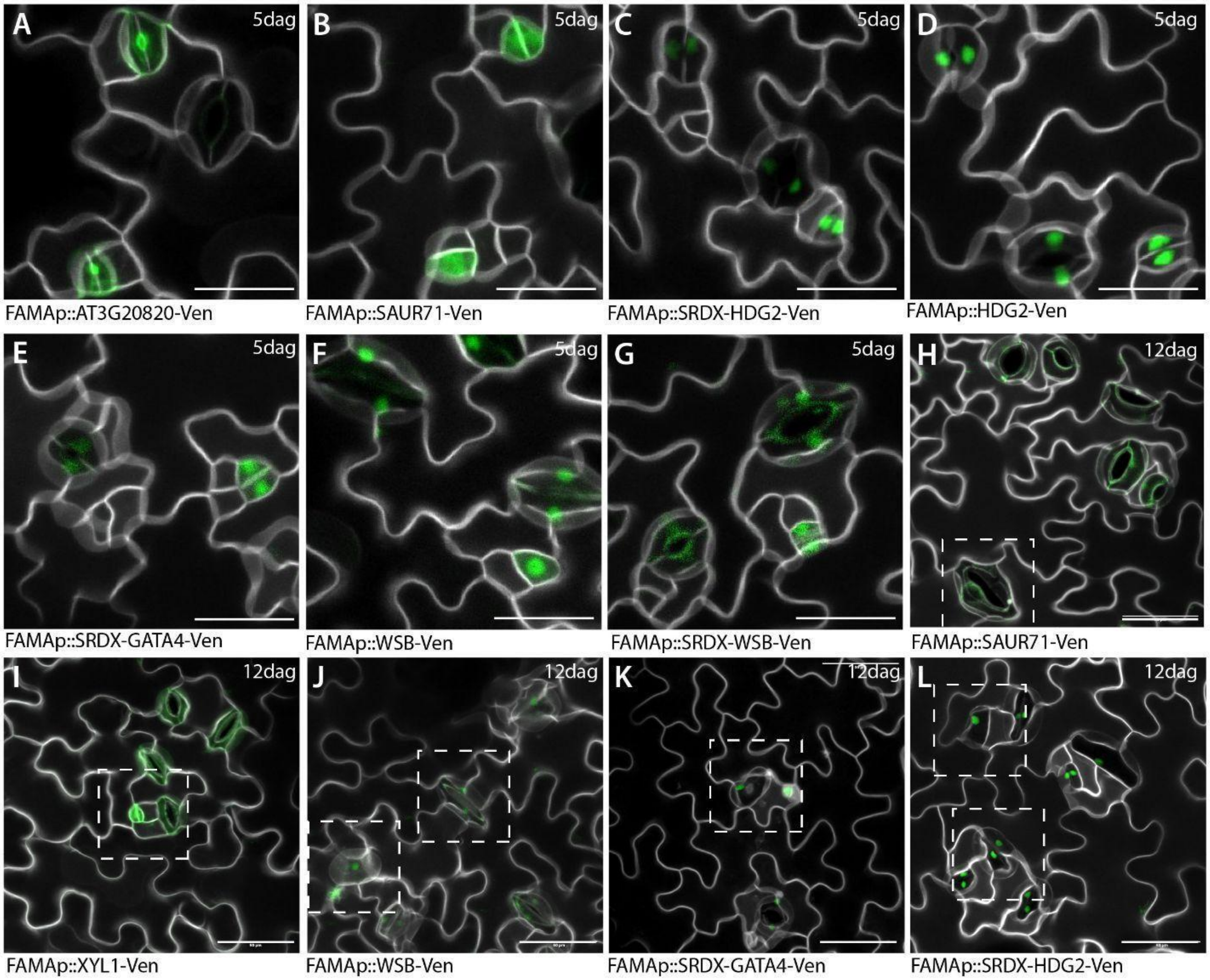


**Figure S11. Subcellular localization of misexpressed putative targets of SPCH and FAMA**

Confocal images of misexpression lines for selected putative targets of SPCH and FAMA in 5 dag (**A-G**) or 12 dag (**H-L**) cotyledons. Dashed squares indicate abnormal stomatal morphology. Scale bars indicate 50 μm. Membranes are visualized using propidium iodide and the plasma membrane marker *ML1p::mCherry-RCl2A* (white).

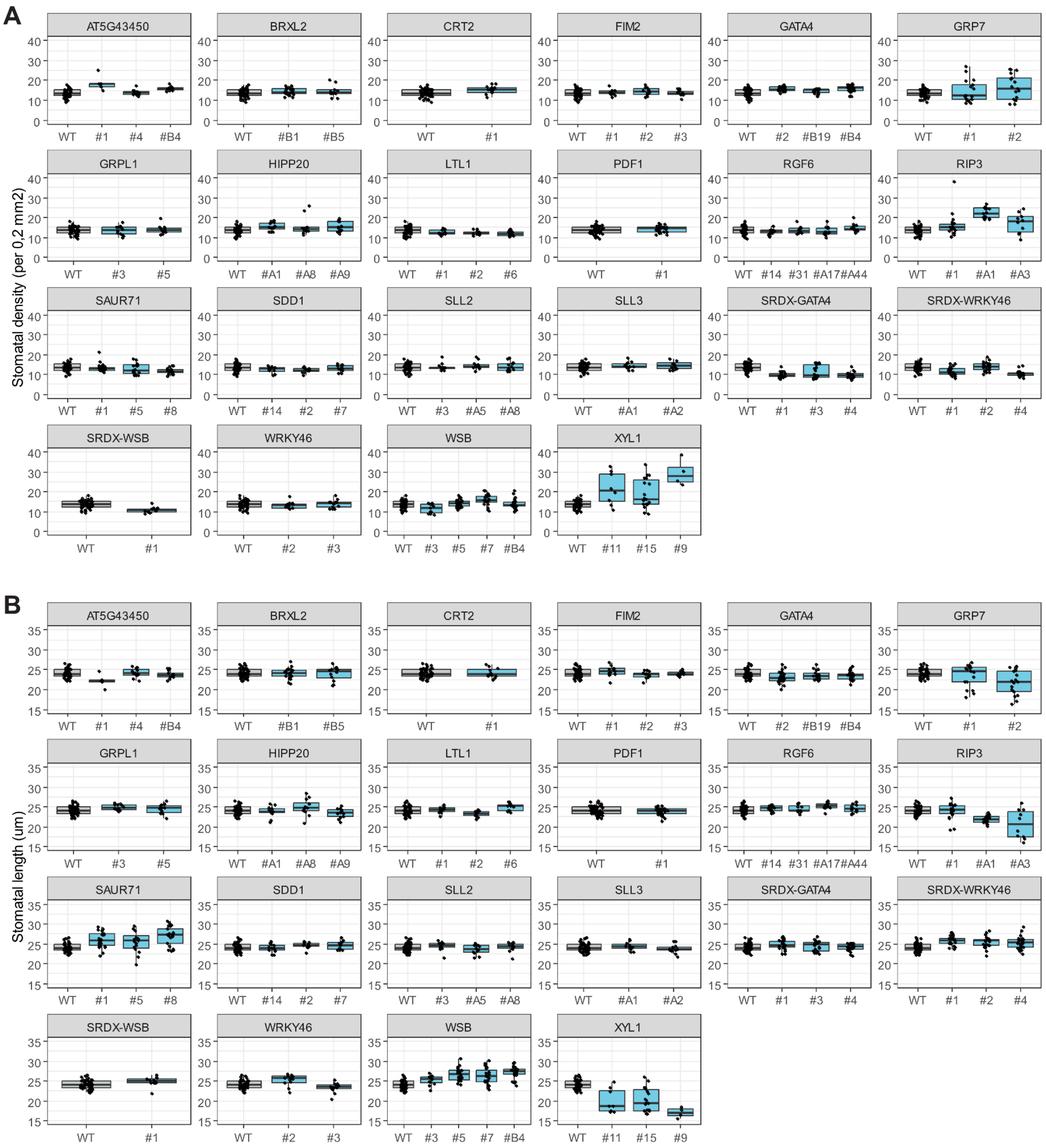


**Figure S12. Changes in stomatal density and size in lines misexpressing putative targets of SPCH and FAMA in the late lineage**

Stomatal density (**A**) and stomatal length (**B**) in 12 dag cotyledons of lines misexpressing putative targets of SPCH and FAMA. Each boxplot corresponds to an independent T2 line.

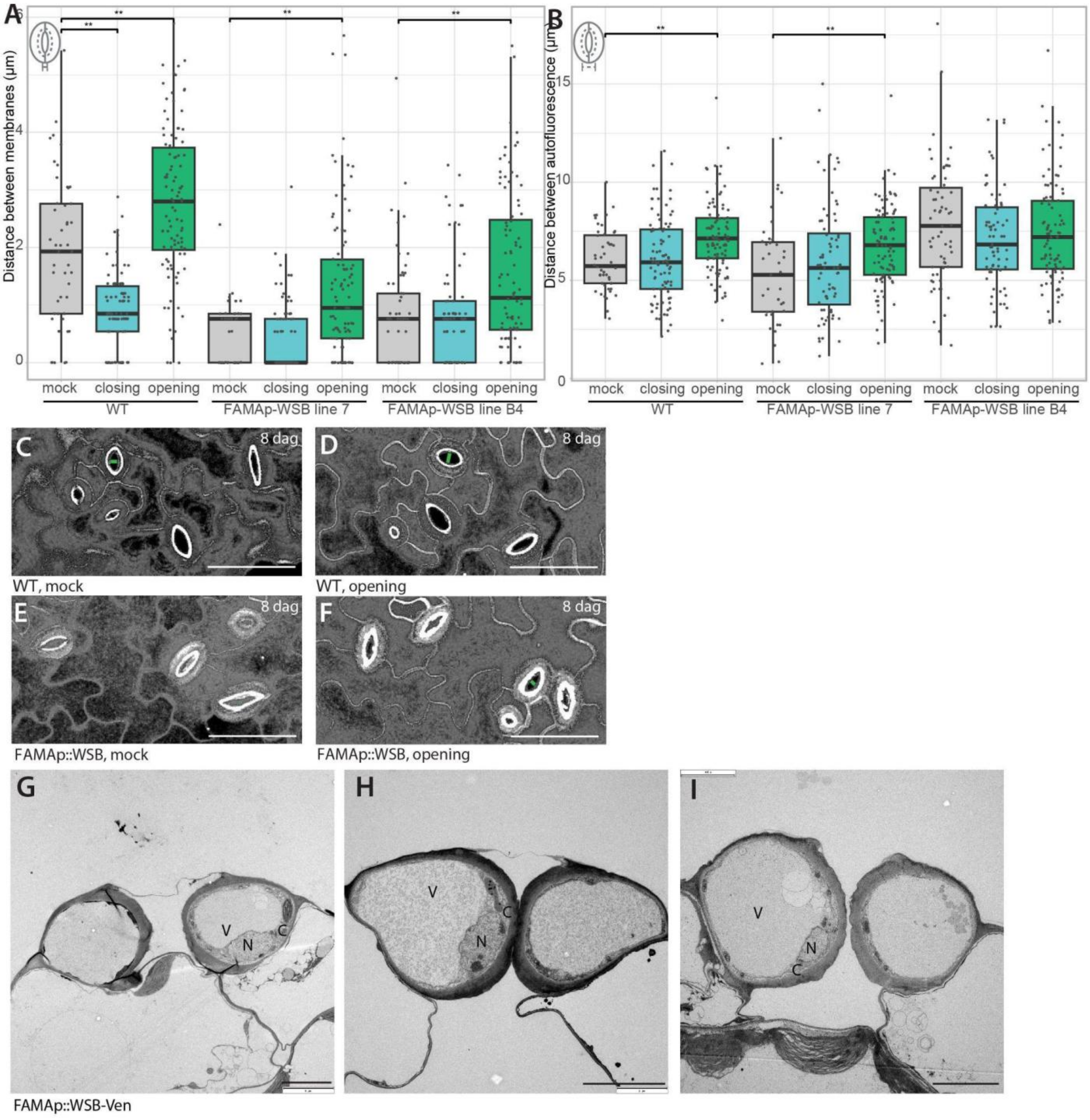


**Figure S13. Misexpression of *WSB* impairs stomatal opening and changes in physical cell morphology**

**A-B.** Longest distance between guard cell membranes (**A**) and between pore autofluorescences (**B**) in 8 dag cotyledons of wild-type and two independent *FAMAp::WSB* lines exposed to closing solution, opening solutions, or mock. Asterisks indicate statistical differences (Student's t test, ** $p<0.01$, *** $p<0.005$)(N=39-98 stomata across 3-5 leaves). **C-F.** Representative confocal z-stacks of wild-type (**C-D**) and *FAMAp::WSB* (**E-F**) exposed to mock or opening solutions. Membranes are visualized using propidium iodide and the plasma membrane marker *ML1p::mCherry-RCl2A* (white). Black areas indicate that no membrane was present throughout the stack. Green lines indicate where distance between guard cell membranes was measured. Scale bars indicate 50 μm. **G-I.** Cross-sectional TEM images showing large guard cells in *FAMAp::WSB*. V =vacuole , N =nucleus , C =cytosol , Scale bars indicate 5 μm.

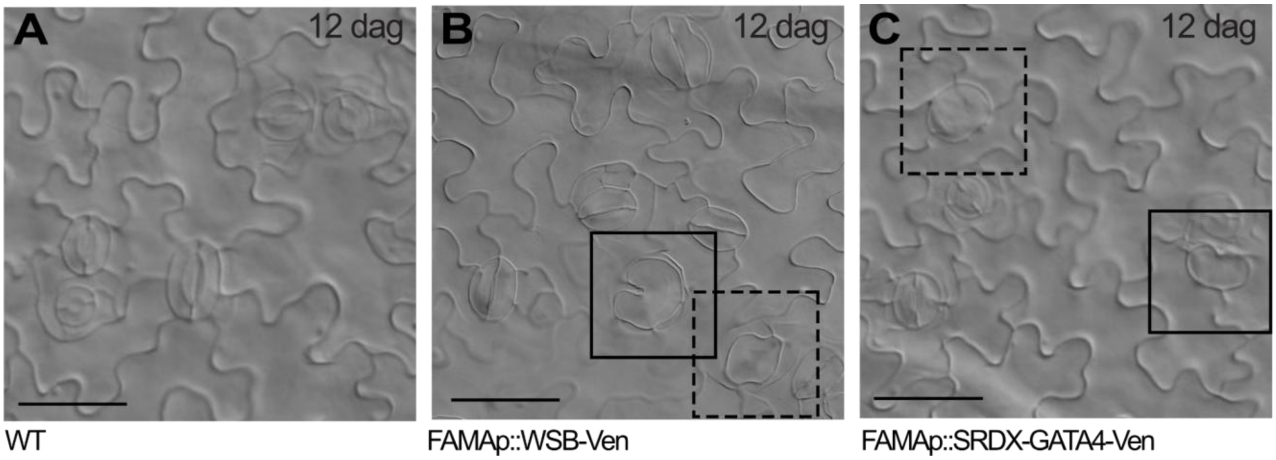


**Figure S14. Misexpression of a subset of putative targets of SPCH and FAMA triggers the formation of single guard cells**

DIC images of 12 dag cotyledons of wild-type (**A**), *FAMAp::WSB-Ven* (**B**) and *FAMAp::SRDX-GATA4-Ven* (**C**). Solid squares highlight large kidney-shaped single GCs and dashed squares indicate large round single GCs. Scale bars indicate 50 µm.

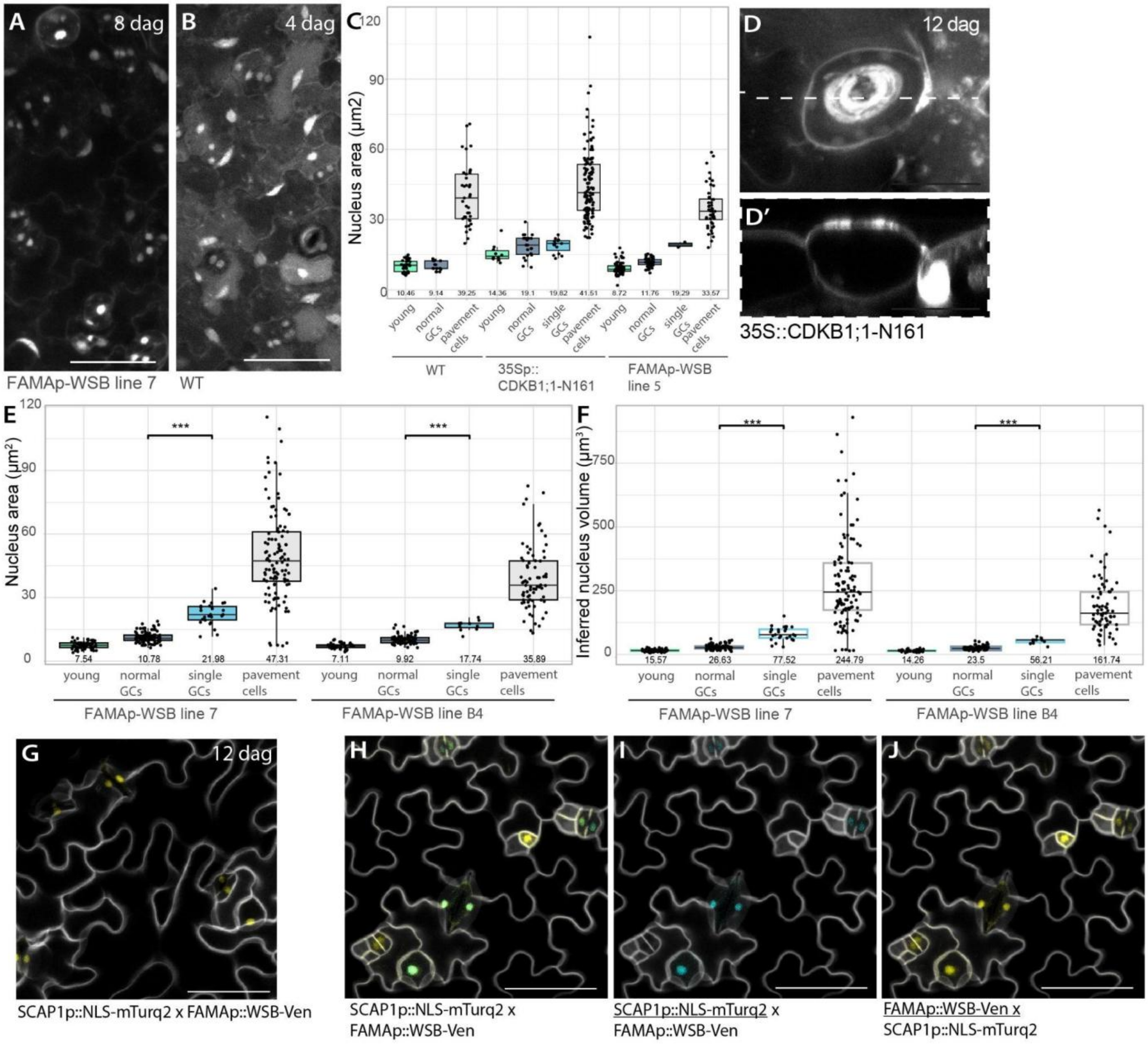


**Figure S15. Single guard cells induced by *WSB* misexpression differentiate after DNA replication**

**A-B.** Confocal z-stacks of Hoechst stained *FAMAp::WSB* and WT leaves used for nucleus area measurements. Scale bars indicate 50 µm. **C.** Graph showing nucleus area of young stomatal cells, regular GCs, SGCs (single guard cells), and pavement cells at 4 dag for WT, *35S::CDKB1;1-N161*, and *FAMAp::WSB*. **D.** Confocal image of large single cells in 12 dag cotyledons of *CDKB1;1-N161*. Scale bars indicate 25 µm. **E.** Additional nucleus area measurements for 2 independent *FAMAp::WSB* lines, expanding in Figure 5E. **F.** Nucleus volume inferred from area measurements in

E. **G.** *FAMAp::WSB* signal for the corresponding image in Figure 5D. **H-I**. Additional images showing SCAP reporter expression in *FAMAp::WSB*. Scale bars indicate 50 μm.

**Table S1. Differentially expressed genes (DEGs) upon FAMA and/or SPCH induction.** The first three columns contain the AGI gene identifier, symbol and synonyms. The following four columns indicate, for each gene, whether it is statistically (FDR-adjusted p-value < 0.05) different between SPCH and FAMA induction, or differentially expressed over time upon FAMA/SPCH induction, and whether the gene has been selected for further analyses (see Results). The following columns contain specific fold-change values for each gene and comparison. A complete description of each gene can be found at the end.

**Table S2. List of DEGs identified that are putative targets of SPCH and/or FAMA.** List containing all DEGs identified in our study (Table S1) that are putative targets of SPCH and/or FAMA as identified by ChIP-seq studies (Lau et al., 2014; Liu et al., 2024).

**Table S3. Annotated table of all DEGs identified in this study.**

**Table S4. Putative targets of SPCH and/or FAMA selected for further analyses.** The first four columns indicate the gene TAIR identifiers, most commonly used symbols and full names, followed by a description. The last three columns indicate whether the gene is differentially expressed in our dataset upon induction of SPCH, FAMA or statistically different between SPCH and FAMA induction.

**Table S5. Cloning primers used in this study**
